# The DBD-α4 helix of EWSR1::FLI1 is required for GGAA microsatellite binding that underlies genome regulation in Ewing sarcoma

**DOI:** 10.1101/2024.01.31.578127

**Authors:** Ariunaa Bayanjargal, Cenny Taslim, Iftekhar A. Showpnil, Julia Selich-Anderson, Jesse C. Crow, Runwei Zhou, Stephen L. Lessnick, Emily R. Theisen

**Affiliations:** Center for Childhood Cancer, Abigail Wexner Research Institute at Nationwide Children’s Hospital, Columbus, OH, 43205, USA; Medical Scientist Training Program, The Ohio State University, Columbus, OH, 43210, USA; Biomedical Sciences Graduate Program, The Ohio State University, Columbus, OH, 43210, USA; Ohio State Biochemistry Program; Department of Pediatrics, The Ohio State Univeristy, Columbus, OH, 43210, USA; Division of Pediatric Heme/Onc/BMT, The Ohio State University College of Medicine, Columbus, OH, 43210, USA

## Abstract

Ewing sarcoma is the second most common bone cancer in children and young adults. In 85% of patients, a translocation between chromosomes 11 and 22 results in a potent fusion oncoprotein, EWSR1::FLI1. EWSR1::FLI1 is the only genetic alteration in an otherwise unaltered genome of Ewing sarcoma tumors. The EWSR1 portion of the protein is an intrinsically disordered domain involved in transcriptional regulation by EWSR1::FLI1. The FLI portion of the fusion contains a DNA binding domain shown to bind core GGAA motifs and GGAA repeats. A small alpha-helix in the DNA binding domain of FLI1, DBD-α4 helix, is critical for the transcription function of EWSR1::FLI1. In this study, we aimed to understand the mechanism by which the DBD-α4 helix promotes transcription, and therefore oncogenic transformation. We utilized a multi-omics approach to assess chromatin organization, active chromatin marks, genome binding, and gene expression in cells expressing EWSR1::FLI1 constructs with and without the DBD-α4 helix. Our studies revealed DBD-α4 helix is crucial for cooperative binding of EWSR1::FLI1 at GGAA microsatellites. This binding underlies many aspects of genome regulation by EWSR1::FLI1 such as formation of TADs, chromatin loops, enhancers and productive transcription hubs.

## Introduction

Ewing sarcoma is an aggressive bone-associated tumor currently treated with dose-intense chemotherapy, radiation, and surgery ***(Pappo and Dirksen, 2018)***. It affects adolescents and young adults with an incidence rate of 3 per million ***(Ozaki, 2015)***. Of these patients, 25-35% have overt metastatic disease with a recurrence rate of 50-80% ***(Gaspar et al., 2015)***. Roughly a quarter of patients with localized disease also relapse ***(Van Mater and Wagner, 2019)***. The five-year survival rate of metastatic and relapsed patients is only 10-30% ***(Stahl et al., 2011)***. Treatment options for relapsed/metastatic Ewing sarcoma patients have not improved for the last four decades. The lack of efficient and targeted treatment for Ewing sarcoma can be attributed to our poor understanding of how precisely Ewing sarcoma is driven by a fusion oncoprotein called EWSR1::FLI1.

Expression of EWSR1::FLI1 results from a translocation between chromosomes 11 and 22 that fuses two genes, *EWSR1* and *FLI1* ***(Delattre et al., 1992)***. As a result, the transcription activation domain of EWSR1 and the DNA binding domain of FLI1 fuse together to create a potent transcription factor ***(May et al., 1993)***. The EWSR1 domain is highly disordered and is important for multimerization and transcriptional hub formation ***(Chong et al., 2018)***. The FLI portion contains a DNA binding domain that binds at two distinct sites: canonical FLI1 binding sites containing a GGAA core and microsatellites containing GGAA repeats ***(Gangwal et al., 2008b)***. At these GGAA microsatellites, EWSR1::FLI1 binding creates de-novo enhancers driving genome-wide reprogram-ming of enhancers to Ewing-specific enhancers ***(Riggi et al., 2014)***. Regulation of Ewing-specific enhancers underlies the mechanisms by which EWSR1::FLI1 regulates expression of thousands of target genes ***(Grunewald et al., 2015)***. More specifically, activation of genes involved in proliferation, migration and invasion pathways leads to oncogenic transformation of precursor cells to Ewing sarcoma cells ***(Kauer et al., 2009)***.

In our recent studies of EWSR1::FLI1, we found a small alpha helix in the DNA binding domain, DBD-α4, to be required for transcription and regulation by the fusion protein ***(Boone et al., 2021)***. Interestingly, this study did not find any change in chromatin accessibility (ATAC-Seq) and genome localization of EWSR1::FLI1 constructs (CUT&RUN) when the DBD-α4 helix was deleted leaving the mechanistic basis for the requirement of the DBD-α4 in transcription regulation unclear. In parallel studies, we also observed that EWSR1::FLI1 expression results in widespread changes to 3D chromatin organization ***(Showpnil et al., 2022)***. These results together prompted us to consider whether the DBD-α4 helix is contributing towards EWSR1::FLI1’s abilities to make changes in the epigenome and the local chromatin structure. To assess whether the DBD-α4 helix is important in chromatin organization, we used our “knock-down/rescue” system in A-673 and TTC-466 cells ***(Theisen et al., 2019; Boone et al., 2021; Showpnil et al., 2022)*** in conjunction with genomics techniques including RNA-Seq, CUT&Tag, and Micro-C. These sets of experiments allowed us to characterize the involvement of the DBD-α4 helix in modulating the chromatin organization, epigenetic reprogramming of enhancers, binding at GGAA microsatellites, and promoting transcription leading to transformation.

## Results

### The DBD-α4 helix of FLI1 domain is implicated in restructuring of 3D chromatin in A-673 cells

To assess the mechanism by which the DBD-α4 promotes transcription, we depleted endogenous EWSR1::FLI1 expression with shRNA (KD) and rescued with previously published DBD and DBD+ EWSR1::FLI1 constructs with 3X-FLAG tags in A-673 cells (Figure 1A, ***Boone et al. (2021))***. The main difference between DBD and DBD+ constructs is the presence of DBD-α4 helix in DBD+ (Supple-mentary Figure 1A). We recapitulated our previous findings that DBD is unable to rescue the same number of Ewing-sarcoma specific genes as DBD+, therefore incapable of driving oncogenic trans-formation (Supplementary Figure 1B-D, ***Boone et al. (2021))***.

**Figure 1.**
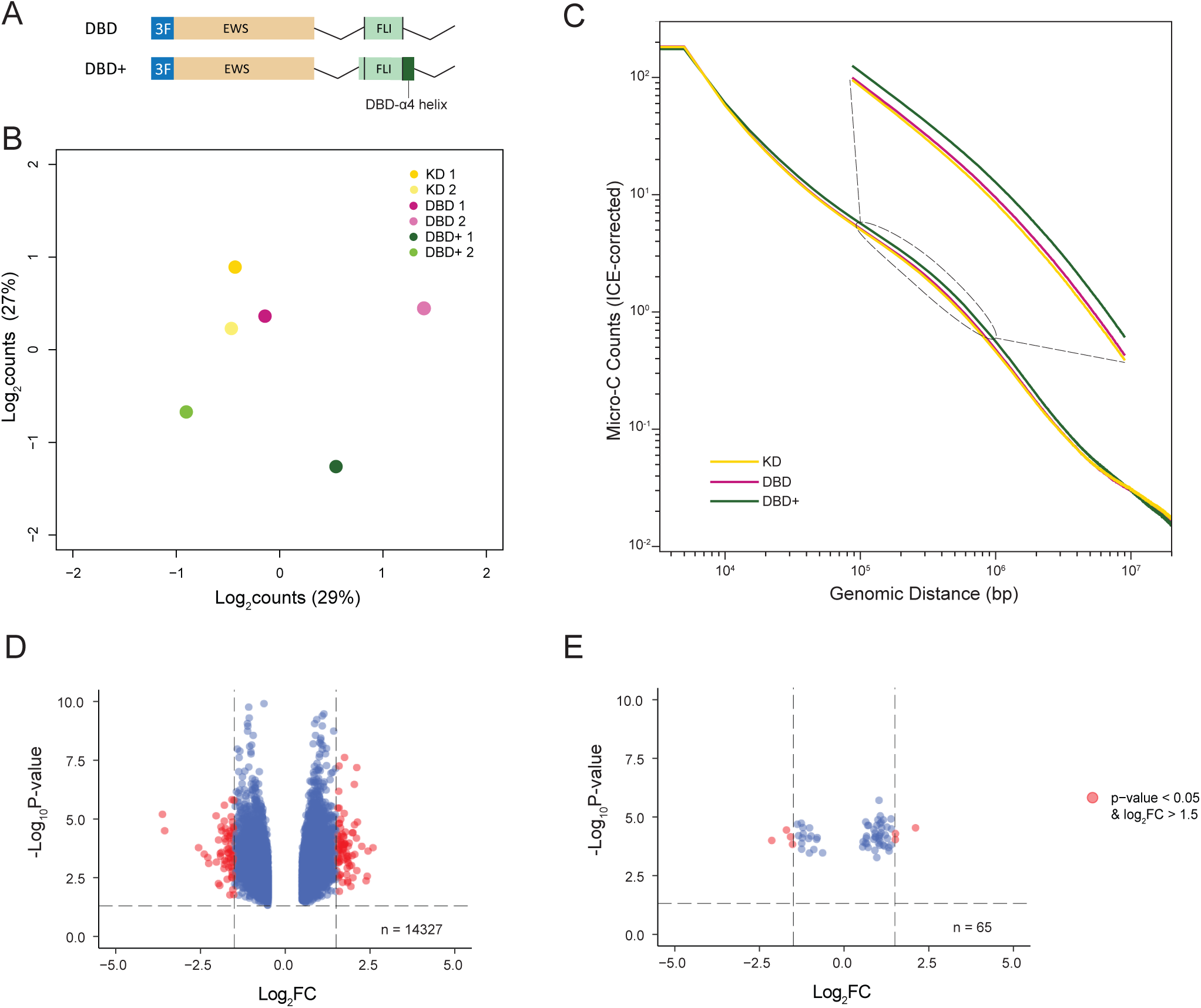
The DBD-*α*4 helix of FLI1 domain is required to restructure chromatin in A-673 cell. A. A schematic of DBD and DBD+ constructs used in shRNA knock-down and rescue experiments. B. Multidimensional scaling (MDS) plot of top 1000 interactions (500kb resolution) in each biological replicates. C. Genome-wide interaction frequency (ICE-corrected Micro-C counts) over genomic distance (bp) at 5kb resolution. D. Volcano plot showing differentially interacting regions (DIRs) detected at 500kb resolution for DBD+ replicates versus KD replicates. E. Volcano plot showing DIRs detected at same resolution for DBD replicates versus KD replicates. Boxplots depict the minimum, first quartile, median, third quartile, and maximum. * P value < 0.05, *** P value < 0.001

Next, we sought to understand the role of DBD-α4 helix in 3D chromatin organization. We carried out Micro-C experiments in KD, DBD and DBD+ cells. First, we performed multidimensional scaling (MDS) analysis of the top 1000 interactions of Micro-C matrices at 500kb resolution to assess the global regulation of 3D chromatin organization (Figure 1B). KD replicates clustered together with DBD replicate 1 on both axes and with DBD replicate 2 on y-axis. DBD+ replicates, on the other hand, clustered away from both KD and DBD replicates. These observations suggest that the global chromatin structure of DBD replicates are more similar to KD than DBD+ replicates. We also assessed the contact frequency across the genome in KD, DBD and DBD+ cells at 5kb resolution (Figure 1C). We found that at the mid region of the plot (10^5^-10^6^ bp), DBD+ cells make more contacts in comparison to KD and DBD cells. This pattern of contact frequency was also detected in replicates 2, but not in replicates 1 when plotted separately (Supplementary 2A-B). At this genome-wide level, we observe that DBD cells display similar contact frequency as KD cells suggesting that the DBD-α4 helix participates in EWSR1::FLI1 mediated 3D chromatin restructuring.

To determine if these changes were significant, we conducted differential interaction analysis using 500kb matrices to compare DBD and DBD+ biological replicates to KD replicates. First, we detected 14,326 and 65 interactions in DBD+ and DBD cells respectively using cut-off values of adjusted p-value < 0.2 and log_2_FC > 0.5. Then, we used cut-off values of adjusted p-value < 0.05 and log_2_FC > 1.5 to identify differentially interacting regions (DIRs). We found that DBD+ cells have 151 DIRs compared to KD cells (Figure 1D) whereas DBD cells only have 7 DIRs (Figure 1E). Taken together, these data demonstrate that the DBD-α4 helix of FLI1 domain is potentially required for EWSR1::FLI1 to restructure 3D chromatin in A-673 cells.

### Altered TAD structure in A-673 DBD cells is linked to GGAA microsatellite binding

To investigate the necessity of the DBD-α4 helix for higher-order chromatin organization, we car-ried out topologically associated domain (TAD) analysis in A-673 cells using HiCExplorer suite of tools ***(Wolff et al., 2020)***. TADs are functional subunits of chromatin that define regulatory land-scape ***(Szabo et al., 2019)***. Within these domains, chromatin interactions including those involving regulatory elements are spatially confined, often by insulators ***(Dixon et al., 2012; Hou et al., 2012; Nora et al., 2012)***. Moreover, genes located in the same TAD are typically co-regulated together ***(Dixon et al., 2016; Ramírez et al., 2018)*** and this co-regulation is facilitated by interactions between enhancers and promoters, which also tend to occur within the confines of a same TAD ***(Bonev et al., 2017)***. Our initial step involved identifying TADs in both DBD and DBD+ matrices with hicFindTADs at 10kb, 25kb, 50kb and 100kb resolutions. Subsequently, we computed differential TADs with hicDifferentialTAD. This analysis entailed a comparison of precomputed TAD regions of DBD and DBD+ matrices with the same regions of KD matrix. The purpose was to detect TADs that appeared specifically in response to the expression of DBD and DBD+ constructs in comparison to KD sample (Figure 2A). At each resolution, the number of TADs detected when DBD+ was expressed was far greater than when DBD was expressed.

**Figure 2.**
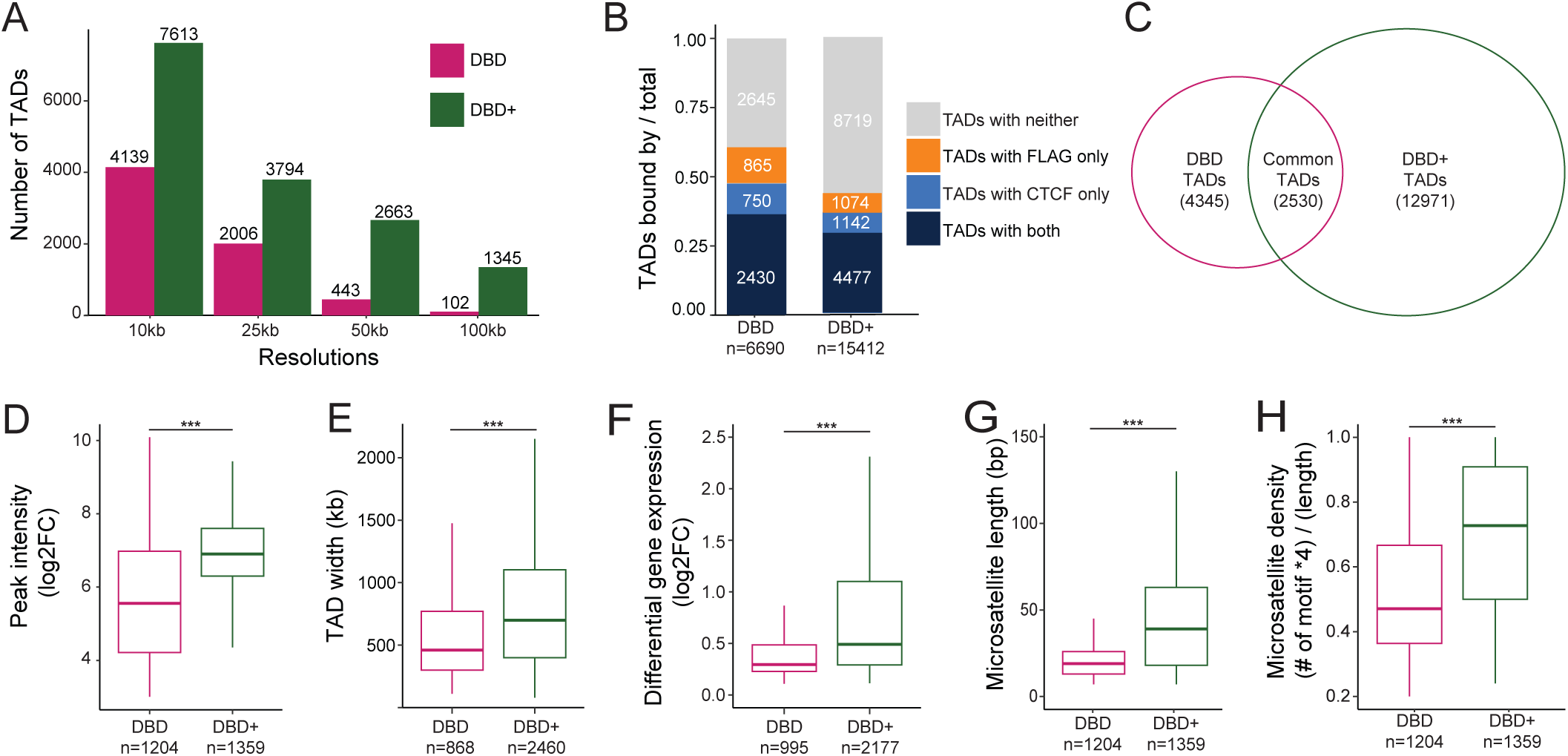
Altered TAD structure in A-673 DBD cells is linked to GGAA microsatellite binding. A. Number of TADs detected in DBD and DBD+ compared to KD at resolutions of 10kb, 25kb, 50kb and 100kb. B. Proportion of TADs (compared to KD) bound by FLAG, CTCF, both, or neither. C. Venn diagram of overlap between DBD and DBD+ TADs (compared to KD). D-H. Comparison of DBD and DBD+ unique TADs. D. Binding intensity of unique FLAG peaks (FDR < 0.05, FC > 8, counts > 80, IDR < 0.01) across the width of DBD and DBD+ unique TADs. E. Width of DBD and DBD+ unique TADs in bp. F. Expression level of significantly upregulated genes within unique TADs in DBD and DBD+ bound by FLAG. G. Length of microsatellites bound by unique FLAG peaks in DBD and DBD+ conditions in bp. H. Density of GGAA motif in the microsatellites calculated as (# of motif x 4)/(length of microsatellites) in DBD and DBD+ unique TADs bound by unique FLAG peaks. Boxplots depict the minimum, first quartile, median, third quartile, and maximum. *** P value < 0.001

To exclude duplicated TADs when combining TADs from different resolutions, we used the default threshold value of 5000bp as the minimum boundary difference required to consider two TADs as distinct. With this approach, we found 6,690 total TADs for DBD and 15,412 total TADs for DBD+ (Figure 2B). To understand the local biological context of these total TADs in DBD and DBD+ cells, we annotated the whole width of the TADs with CTCF and FLAG CUT&Tag peaks (FDR < 0.05, FC >8, counts > 80, IDR < 0.01) (Figure 2B). In DBD cells, we detected 11.2% of the TADs bound by CTCF only, 12.9% bound by DBD FLAG only, 36.3% bound by both CTCF and DBD FLAG, and 39.6% bound by neither. In contrast, for DBD+, we detected 7.4% and 7.0% of the TADs bound by only CTCF and only DBD+ FLAG, respectively, 29.0% bound by both and 56.6% bound by neither. These observations highlight both the increased number of total TADs detected in DBD+ cells and a notable increase in the percentage of TADs detected in DBD+ cells with no CTCF or FLAG association. This suggests a broader role for the DBD-α4 helix in genome-wide reorganization of TADs by EWSR1::FLI1, consistent with the results discussed above relating to Figure 1.

Next, we overlapped total TADs from DBD and DBD+ conditions and identified 2,530 TADs common to both. There were 4,345 unique TADs in DBD cells and 12,971 unique TADs in DBD+ cells (Figure 2C). To understand the transcriptional activity driven by the DBD-α4 helix at GGAA repeats, we focused on the whole width of unique TADs containing DBD and DBD+ FLAG peaks at GGAA microsatellites (Figure 2D-2H). This analysis narrowed the TADs down to 868 unique ones for DBD cells and 2,460 unique TADs for DBD+ cells (Figure 2E). We observed that within these TAD sub-sets containing FLAG peaks at GGAA microsatellites, the intensity of the DBD+ FLAG peaks was higher compared to DBD FLAG peaks (Figure 2D). Moreover, TADs uniquely bound by DBD+ at GGAA microsatellites were not only wider (Figure 2E) but also contained a higher number of differentially expressed genes compared to those bound only by DBD (Figure 2F and Supplementary Figure 3A). Further analysis of GGAA microsatellites bound by DBD and DBD+ peaks within unique TADs revealed that the DBD+ FLAG peaks were associated with longer microsatellites (Figure 2G) and denser GGAA motifs (Figure 2H, density calculated by multiplying the number of GGAA motif by 4 and dividing by the total length of the microsatellites) in contrast those bound by DBD alone. These findings suggest that DBD+ accesses longer and denser microsatellites within larger TADs and binds to these sites with higher binding intensity than DBD. Furthermore, these TADs with DBD+ bound microsatellites show greater changes in genes expression suggesting a functional role for the DBD-α4 helix in gene regulation within TADs.

TAD boundaries are regions between TADs that act as insulators ***(Dixon et al., 2012)*** and are often marked by active chromatin ***(Ay et al., 2014)***. We conducted a similar analysis as above in a 20kb region around unique TAD boundaries (Supplementary Figure 3B-3G). We observed a comparable percentage of the unique TAD boundaries were bound by only CTCF peaks in DBD and DBD+ cells (36.3% and 32.2%, respectively). The percentage of boundaries with FLAG peaks in DBD and DBD+ cells were 15.2% and 8.6%, respectively (Supplementary Figure 3B). When we considered boundaries that overlap with FLAG peaks at microsatellites (Supplementary Figure 3C-3G), we observe similar findings as those spanning the whole width of TADs (Figure 2D, 2F-2H). The FLAG peaks at DBD+ specific TAD boundaries were of higher intensity than FLAG peaks at DBD specific TAD boundaries (Supplementary Figure 3C). Next we assessed gene expression at TAD boundaries and found DBD+ boundaries contained more significantly regulated genes (Supplementary Figure 3D-3E). The GGAA microsatellites bound by DBD+ FLAG peaks at boundaries were also longer and denser than DBD bound microsatellites (Supplementary Figure 3F-3G). These data demonstrate that the DBD-α4 helix affects overall DNA binding at GGAA microsatellites and gene expression in the context of TAD organization.

**Figure 3.**
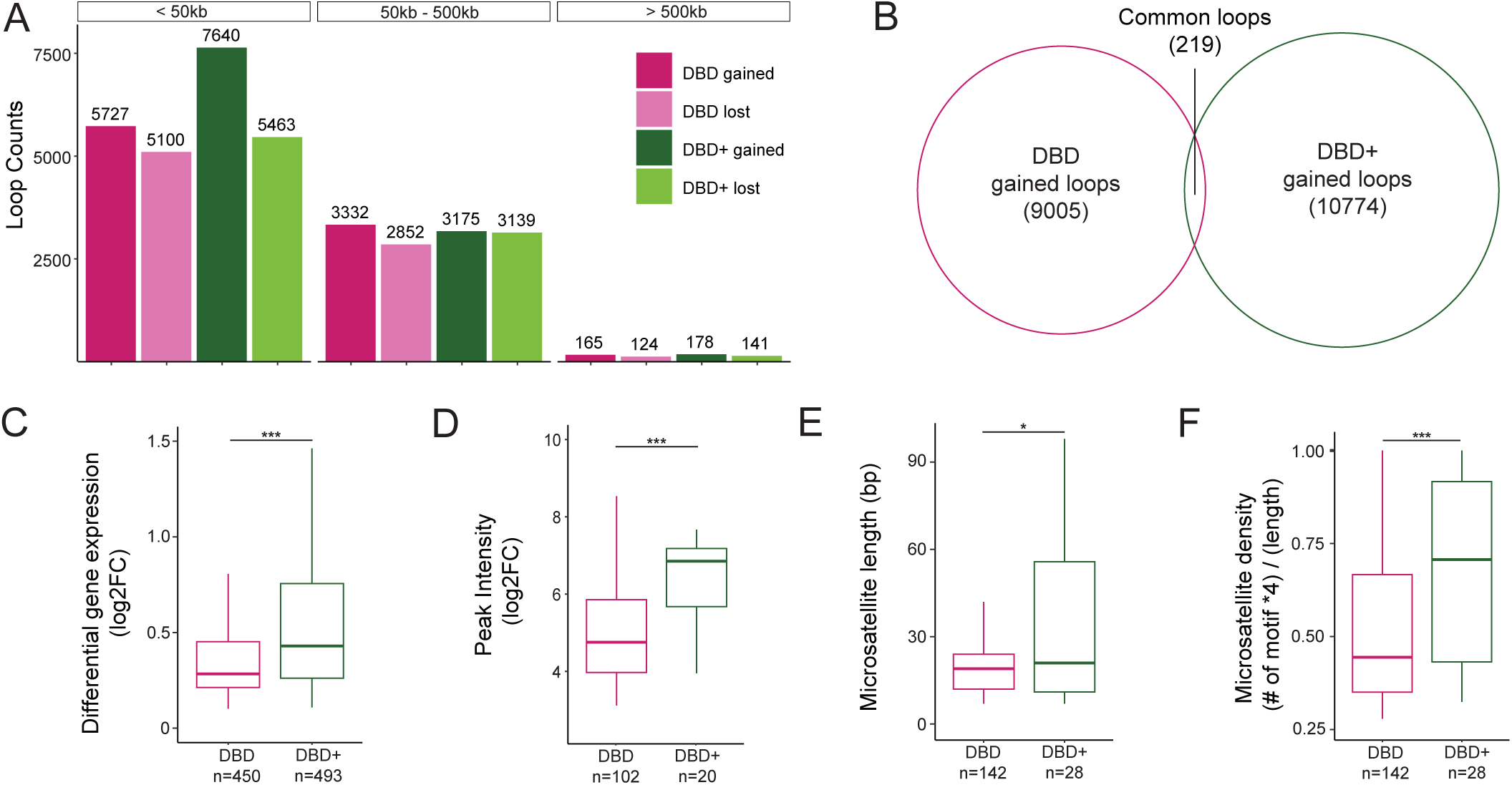
DBD and DBD+ form loops at GGAA microsatellites, but DBD+ rescues more and shorter loops in A-673 cells. A. Number of loops detected in DBD and DBD+ compared to KD at resolutions of 1kb at short-range (<50kb), mid-range (50kb-500kb), and long-range (>500kb). B. Venn diagram of overlap between DBD and DBD+ uniquely gained loops (compared to KD). C. Expression level of significant genes overlapped with uniquely gained loop anchors of DBD and DBD+. Means = 0.45 (DBD) and 0.77 (DBD+) D. Peak intensity of unique FLAG peaks (FDR < 0.05, FC > 8, counts > 80, IDR < 0.01) at anchors of uniquely gained loops in DBD and DBD+ cells. Means = 5.01 and 6.30 per DBD and DBD+ E. Length of microsatellites bound by unique FLAG peaks at the anchors of DBD and DBD+ uniquely gained loops in bp. Means = 22.5bp (DBD) and 38bp (DBD+) F. Density of GGAA motif in the microsatellites calculated as (# of motif x 4)/(length of microsatellites) at the anchors of DBD and DBD+ uniquely gained loops bound by unique FLAG peaks. Means = 0.51 (DBD) and 0.70 (DBD+). Boxplots depict the minimum, first quartile, median, third quartile, and maximum.* P value < 0.05, *** P value < 0.001

### DBD and DBD+ form loops at GGAA microsatellites, but DBD+ rescues more and shorter loops in A-673 cells

Previously, we observed that EWSR1::FLI1 promotes chromatin loop formation at GGAA microsatellites leading to enhancer activation and upregulation of gene expression ***(Showpnil et al., 2022)***. We next asked whether the DBD-α4 helix participates in functions of EWSR1::FLI1 pertaining to the formation of loops by calling chromatin loops with Mustache algorithm ***(Roayaei Ardakany et al., 2020)*** in matrices of A-673 DBD and DBD+ cells compared to KD cells at 1kb resolution. We detected loops in 3 ranges defined as short-range (<50kb), mid-range (50kb-500kb), and long-range (>500kb) loops based on the linear genomic distance between loop anchors (Figure 3A). The only range with a difference between DBD and DBD+ cells was short-range loops. Specifically, we found that DBD+ expressed cells gained 1,913 more short-range loops than DBD expressed cells compared to KD. Both conditions had a similar number of short-range lost loops (5,100 and 5,463 for DBD and DBD+ respectively, Figure 3A). For an overlap analysis between DBD and DBD+ conditions, we included all lost loops and found 5610 loops that were commonly lost in both conditions (Supplementary Figure 4A). When the same overlap analysis was carried out for DBD and DBD+ gained loops of all ranges, we found only 219 common loops (Figure 3B). This small overlap of gained loops suggested that although they have a similar number of total gained loops, DBD and DBD+ cells contain loops with very different anchors. These sets of observations suggest that formation of distinct set of short-range loops is a potential mechanism by which the DBD-α4 helix restructures chromatin organization in A-673 cells.

Prompted by the above difference in short-range loop formation, we further investigated uniquely gained loops of all ranges in DBD (9,005) and DBD+ (10,774) conditions (Figure 3B). When individual anchors of loops were overlapped with GGAA microsatellites, we found 92.4% and 92.2% of DBD and DBD+ loop anchors were at GGAA microsatellites. We then assessed overall gene expression at all uniquely gained loop anchors and found DBD+ loop anchors associated with more gene activation (Figure 3C) and downregulation compared to DBD specific anchors (Supplementary Figure 4B). Next, we focused our analysis on gained loop anchors with FLAG peak at GGAA microsatellites in DBD and DBD+ cells. We found 102 and 20 FLAG peaks overlapping with DBD and DBD+ loop anchors respectively (Figure 3D). At this subset, FLAG peaks had a higher intensity in DBD+ compared to DBD expressed cells. The GGAA microsatellites overlapping with DBD and DBD+ FLAG peaks at gained loop anchors were further characterized in length and density (Figure 3E-F). We found that DBD+ FLAG peaks were bound at GGAA microsatellites that are long and dense in comparison to DBD bound microsatellites. Taken together, these data indicate that the DBD-α4 helix facilitates formation of short-range loops by promoting binding at long and dense GGAA microsatellites, thus leading to more gene regulation.

### DBD+ rescues de-novo enhancer formation at GGAA microsatellites in A-673 cells

Multiple studies have shown that binding of EWSR1::FLI1 at GGAA microsatellites results in genome-wide de-novo enhancer formation ***(Riggi et al., 2014; Tomazou et al., 2015; Sheffield et al., 2017)***. Ewing-specific enhancer establishment at GGAA microsatellites are functionally linked to the onco-genic transformation ***(Johnson et al., 2017a)***. We therefore sought to understand whether the DBD-α4 helix of EWSR1::FLI1 has any role in modulating the Ewing-specific enhancer landscape. To assay the enhancer landscape, we collected H3K27ac CUT&Tag data from A-673 KD, DBD and DBD+ cells. Principal component analysis of H3K27ac localization shows that the DBD replicates were clustered closer to the KD replicates while being in between the KD and the DBD+ replicates (Figure 4A), suggesting that the DBD-α4 helix is required to reshape the enhancer landscape. Next, we did overlap analysis of H3K27ac peaks of DBD and DBD+ cells. There were 7,854 H3K27ac peaks unique to DBD cells, 35,583 peaks common between DBD and DBD+ cells, and 26,317 peaks unique to DBD+ cells (Figure 4B), highlighting the vastly different enhancer landscapes of DBD and DBD+ conditions. Next, we sought to understand the extent of GGAA microsatellite driven H3K27 acetylation by conducting overlap analysis of H3K27ac peaks and GGAA microsatellites that were previously characterized (Figure 4C, ***Johnson et al. (2017a))***. Interestingly, for each group of H3K27ac peaks we considered (i.e. DBD, common, and DBD+), we found around 40% of the H3K27ac peaks overlapped with GGAA microsatellites. Overall, a significant portion of H3K27ac peaks correlated with GGAA microsatellites in both DBD and DBD+ conditions underscoring the importance of GGAA repeats in formation of novel enhancers by fusions involving the transcriptional activation domain of EWSR1 with the FLI1 DNA binding domain. However, these results suggest that the DBD-α4 helix is required for modulating the enhancer landscape in Ewing cells to specifically promote oncogenic transformation.

**Figure 4.**
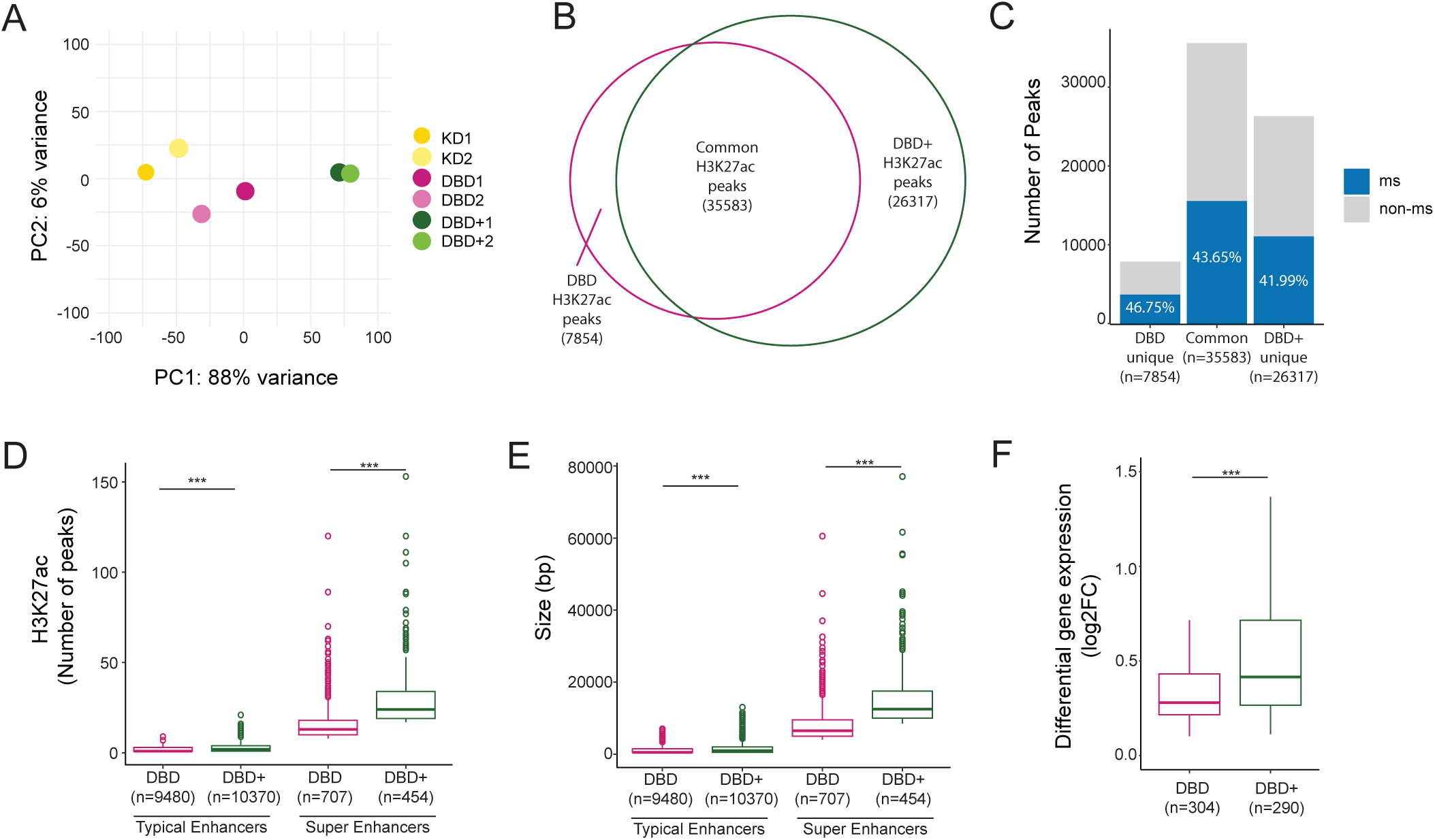
DBD+ rescues de-novo enhancer formation at microsatellites A-673 cells. A. PCA plot of H3K27ac peaks in biological replicates of KD, DBD and DBD+. B. Venn diagram of overlap of H3K27ac peaks (FDR < 0.05, FC > 8, counts > 80, IDR < 0.01) of DBD and DBD+. C. Percentage of H3K27ac peaks at microsatellites in common, DBD unique and DBD+ unique peaks. D. Number of H3K27ac peaks constituting typical and super enhancers called in DBD and DBD+ conditions. E. Constituent size (in bp) of typical and super enhancers in DBD and DBD+ conditions. F. Expression level of significantly upregulated genes at DBD and DBD+ super enhancers. Boxplots depict the minimum, first quartile, median, third quartile, and maximum. Circles depict outliers. *** P value < 0.001.

Transcription factors have been shown to establish super-enhancers at key genes that drive cellular identity ***(Whyte et al., 2013)***. We therefore sought to understand whether the DBD-α4 helix participates in formation of super-enhancers. We utilized ROSE algorithm to define enhancers and super-enhancers in DBD and DBD+ cells ***(Lovén et al., 2013; Whyte et al., 2013)***. We found 9,480 and 10,370 typical enhancers in DBD and DBD+, respectively (Figure 4D). We also identified 707 and 454 super-enhancers in DBD and DBD+, respectively. Although the super-enhancers associated with DBD+ were fewer in numbers compared to DBD, these super-enhancers were larger in size (Figure 4E) and contained more H3K27ac peaks (Figure 4D). We also assessed the expression level of genes associated with super-enhancers and found DBD+ super-enhancer genes had more gene activation (Figure 4F) and deactivation (Supplementary Figure 5A). These data suggest the DBD-α4 helix is required in formation of enhancers and super-enhancers, therefore important in regulation of genes from those highly clustered regions of the genome.

### The DBD-α4 helix promotes binding at longer and denser GGAA microsatellites in A-673 cells

The results of TAD, loop and enhancer analysis hinted towards a mechanism of the DBD-α4 helix in binding GGAA microsatellites that differed from our prior CUT&RUN analysis which showed no difference in binding ***(Boone et al., 2021)***. We decided to further investigate binding patterns of DBD and DBD+ FLAG peaks genome-wide from our A-673 CUT&Tag dataset generated for this study. We found 8,118 DBD+ and 10,101 DBD FLAG peaks genome-wide that passed the threshold of FDR < 0.05, FC > 8, counts > 80, IDR < 0.01. When we overlapped the two sets of FLAG peaks, we found 6,102 common, 3,999 DBD unique and 2,016 DBD+ unique FLAG peaks (Figure 5A). When each group of FLAG peaks was overlapped with genomic GGAA microsatellites, we found that 70.68% of DBD+ unique and 73.04% of common peaks bind at GGAA microsatellites (Figure 5B). When we considered DBD unique peaks, we found GGAA microsatellites binding shifted down to 53.04% (Figure 5B, DBD bar). Common FLAG peaks were found to have the highest peak intensity, followed by DBD+ peaks then DBD peaks (Figure 5C). Taken together, these data demonstrate a role of the DBD-α4 helix in binding at a subset of GGAA microsatellites.

**Figure 5.**
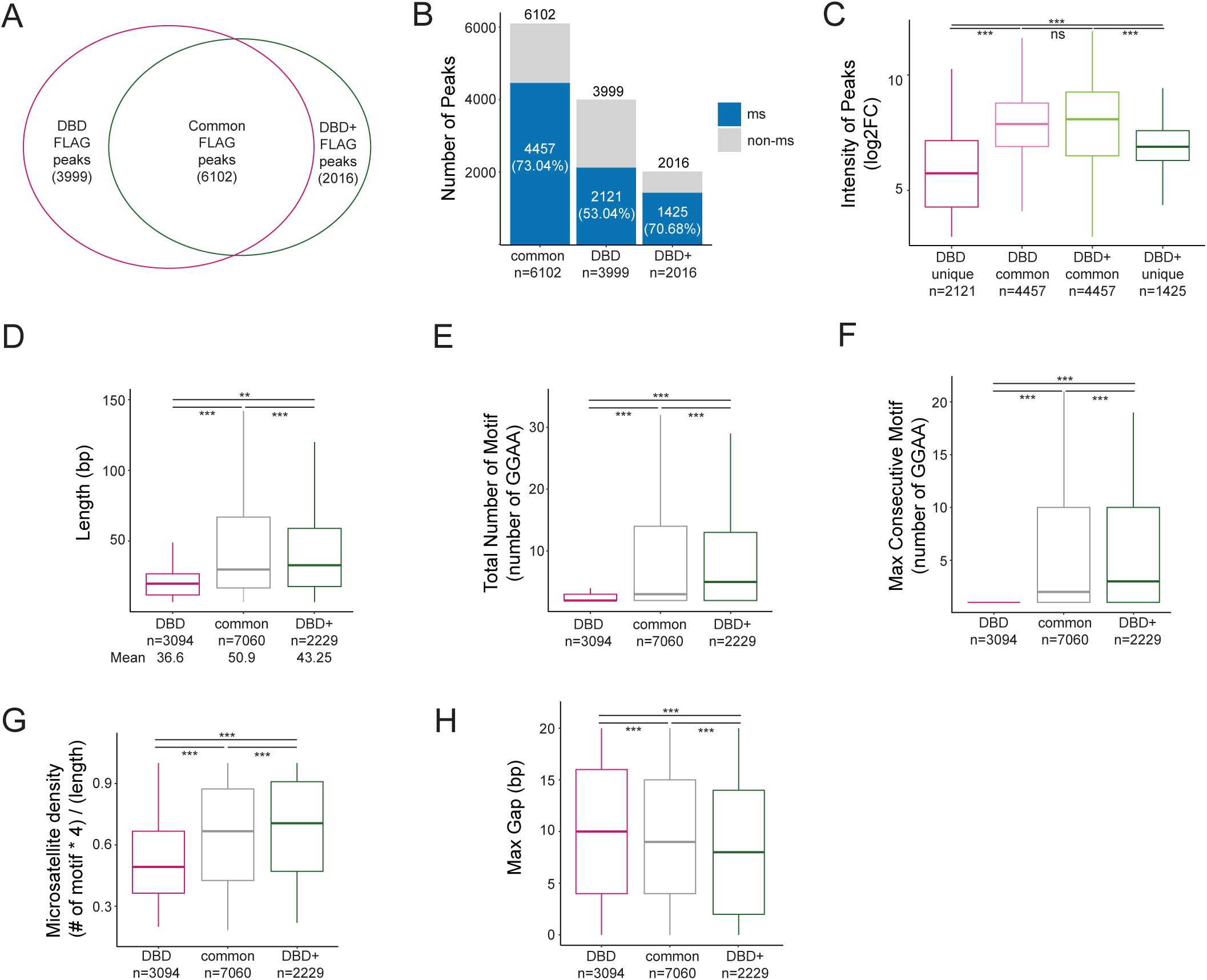
DBD-*α*4 helix of FLI1 promotes binding at longer and denser GGAA microsatellites in A-673 cells. A. Venn diagram of overlap between FLAG peaks (FDR < 0.05, FC > 8, counts > 80, IDR < 0.01) of DBD and DBD+ cells. B. Percentage of FLAG peaks bound at microsatellites in common, DBD unique and DBD+ unique peaks. C. Intensity of peaks in DBD unique (mean=5.75), DBD common (mean=7.76), DBD+ common (mean=7.75) and DBD+ unique (mean=6.83) FLAG peaks. D. Length (in bp) of GGAA microsatellites bound by DBD unique (mean=36.61), common in both (mean=50.94), and DBD+ unique (mean=43.25) FLAG peaks. E. Total number of GGAA motifs in microsatellites bound by DBD unique (mean=4.66), common in both (mean=8.70), and DBD+ unique (mean=7.78) FLAG peaks. F. Maximum consecutive number of GGAA motifs in microsatellites bound by DBD unique (mean=1.48), common in both (mean=5.21), and DBD+ unique (mean=5.32) FLAG peaks G. Percent of GGAA motif in the microsatellites calculated as (# of motif x 4)/(length of microsatellites) bound by DBD unique (mean=0.54), common in both (mean=0.65), and DBD+ unique (mean=0.69) FLAG peaks. H. Maximum number of insertion (gaps in bp) in microsatellites bounds by DBD unique (mean=10.1), common (mean=9.1), and DBD+ unique (mean=8.3) FLAG peaks. Boxplots depict the minimum, first quartile, median, third quartile, and maximum. * P value < 0.05, ** P value < 0.01, and *** P value < 0.001

We therefore further characterized the GGAA microsatellites bound by these 3 sets of peaks: DBD unique, common and DBD+ unique peaks (Figure 5D-H). We assessed the length of microsatellites and found uniquely DBD bound GGAA microsatellites to be shorter (mean=36.61bp) than uniquely DBD+ bound GGAA microsatellites (43.25bp) (Figure 5D). GGAA microsatellites bound by both DBD and DBD+ were the longest (50.94bp) (Figure 5D). When we quantified the total number of GGAA repeats, we found that the mean number of GGAA motifs in uniquely DBD bound GGAA microsatellites was 4.66 compared to 7.78 in uniquely DBD+ bound GGAA microsatellites and 8.70 in GGAA microsatellites commonly bound in both conditions that were significantly different from each other (Figure 5E). Next, we quantified the maximum consecutive GGAA repeats within a microsatellite and found that the DBD unique peaks contained GGAA repeats no longer than 2 repeats (Figure 5F) whereas common and DBD+ unique peaks were localized at approximately 5 consecutive GGAA repeats (mean = 5.21 and 5.32 for common and DBD+, respectively, Figure 5F). Next we plotted GGAA motif density and observed that DBD unique microsatellites are on average (mean=0.54) less dense compared to common (0.65) and DBD+ (0.69) bound GGAA microsatellites (Figure 5G). Inverse of GGAA density is the max gap or number of non-GGAA bp in the microsatellites. We then assessed the gap size in each of the groups and found DBD+ binds microsatellites that contain the least amount of gaps (mean=8.3) followed by common (9.1) then DBD unique peak microsatellites (10.1) (Figure 5H). These analyses demonstrate that the DBD-α4 helix is required for effective binding at GGAA microsatellites that are medium in length and denser in its GGAA motifs.

### The DBD-**α**4 helix is necessary for productive transcription hub formation at GGAA microsatellites in A-673 cells

Transcription is a process that is regulated across large genomic distances, time, and space ***(Cramer, 2019; Lee and Young, 2013)***. One of the emerging concepts that unifies observations of such a regulation is the concept of transcriptional hubs. Transcriptional hubs are actively transcribed regions containing clusters of transcription factors and RNA Pol II, and they are highly characterized by enhancers ***(Lim and Levine, 2021)***. EWSR1::FLI1 has been shown to form such hubs via the intrinsically disordered region of EWSR1 at GGAA microsatellites ***(Chong et al., 2018)***. Since the DBD-α4 helix is required in effective binding at GGAA microsatellites and the downstream regulation of 3D chromatin, we sought to understand if the DBD-α4 helix participates in the function of EWSR1::FLI1 in the formation of transcriptional hubs.

To characterize EWSR1::FLI1-driven transcription hubs, we turned to a region containing the *FCGRT* gene in A-673 cells since EWSR1::FLI1 at this locus is shown to form a hub at GGAA repeats (Figure 6A, ***Gangwal et al. (2008a); Chong et al. (2018)***). The *FCGRT* gene resides in a 250kb TAD (top panel) in both DBD and DBD+ cells. Considering that TADs are highly self-interacting regions and that interactions across different TADs are limited by insulator elements ***(Dixon et al., 2012; Hou et al., 2012; Nora et al., 2012)***, we used TADs as proxy for hubs as we did not expect hubs to extend beyond TAD boundaries. Within the hub, we observed instances where DBD and DBD+ clusters appear in similar locations. In DBD cells, however, we noticed increases in the intensity of all peaks except the *FCGRT* promoter GGAA microsatellite peak (Figure 6A, magenta tracks). When we zoomed in on this microsatellite, DBD and DBD+ peaks had a similar binding pattern (Figure 6B, magenta and green tracks). Despite this similarity, DBD binding did not facilitate the formation of chromatin loops at the *FCGRT* promoter as efficiently as DBD+ (inverted red arcs). Furthermore, this lack of loop formation correlated with decreased enhancer marks (Figure 6A, green tracks and bar) and decreased expression of the *FCGRT* gene (black tracks, padj < 0.05, FC > 2). Even though binding at the *FCGRT* microsatellite is comparable between DBD and DBD+, DBD is less able to modulate transcription and chromatin at this microsatellite with high GGAA motif density (0.79, Figure 6B).

**Figure 6.**
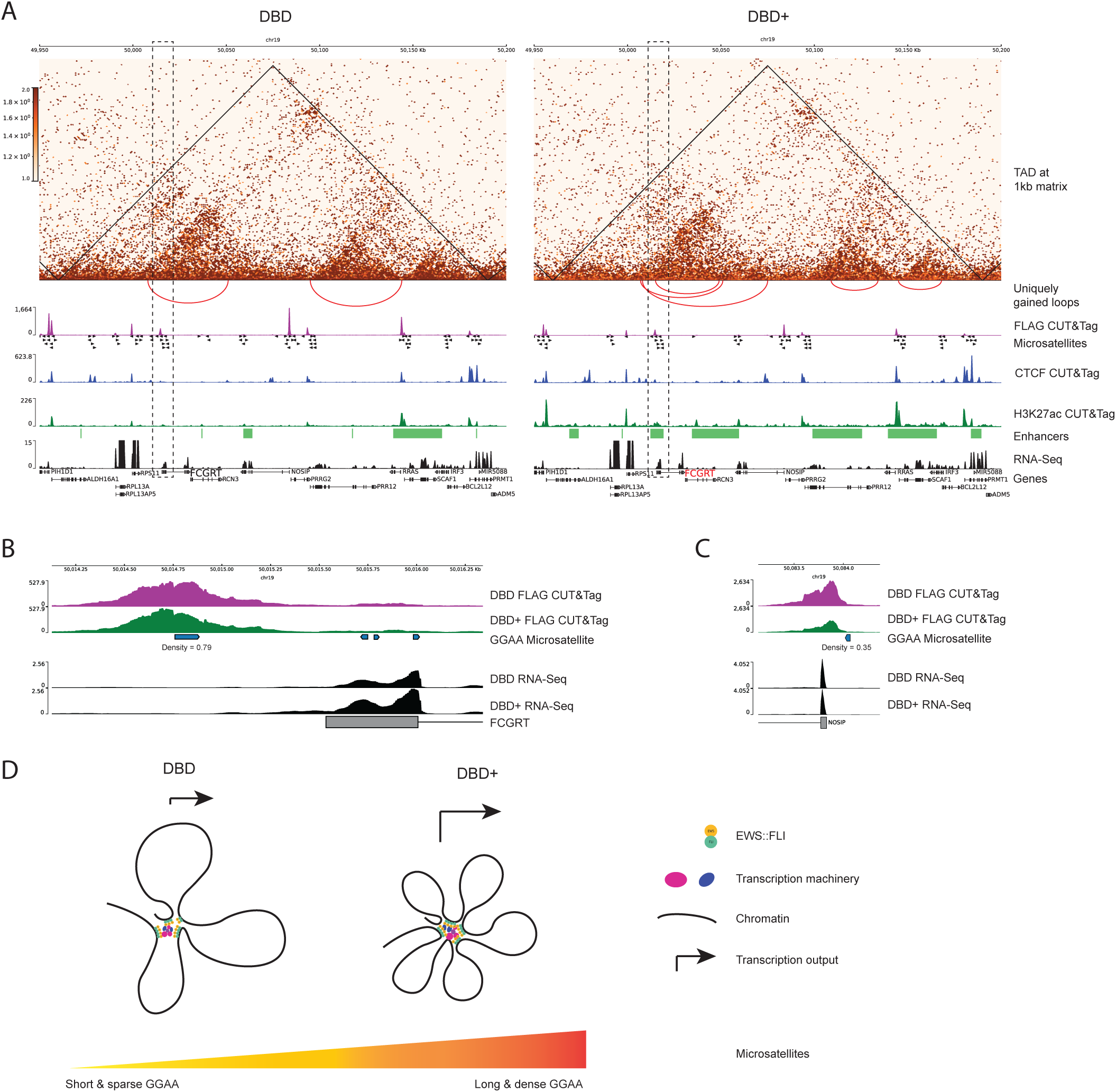
The DBD-*α*4 helix promotes formation of transcription hubs by effective binding at microsatellites. A. 250kb region on chr 19 containing FCGRT and other genes. TADs are depicted on 1kb matrices (DBD/KD and DBD+/KD). Uniquely gained loops are shown as red inverted arcs. FLAG CUT&Tag bigwig tracks depicted in magenta. GGAA microsatellites in hg19. CTCF CUT&Tag track is in blue middle row. H3K27ac tracks are in green. Enhancers and super-enhancers are shown as green bars. Gene expression is in black tracks. B. FCGRT promoter region containing GGAA microsatellites. C. NOSIP promoter region containing GGAA repeats. D. Model of EWSR1::FLI1-driven transcription hub

We also studied the DBD peaks that had higher intensity compared to DBD+ peaks. The majority of these peaks were at microsatellites of varying density (0.35-0.75) and two peaks at the promoters of *PRR12* and *RRAS* were not at GGAA repeats (Figure 6A). These peaks were still near GGAA microsatellites with densities of 0.53 and 0.28 for *PRR12* and *RRAS* peaks respectively. Upon analysis of the RNA-sequencing data, we found that the expression of genes associated with this subset of peaks were not significantly differentially expressed in both DBD and DBD+ cells compared to KD (Supplementary Table 1). The only gene with significantly changed expression was *FCGRT* and only in DBD+ cells compared to KD. Taken together, these data demonstrate that the DBD-α4 helix is required in regulating expression of the *FCGRT* gene.

Another locus we investigated in A-673 cells is *CCND1*, where hub formation at GGAA repeats is also reported ***(Riggi et al., 2014; Chong et al., 2018)***. The *CCND1* locus is located at the border of a 700kb TAD in both DBD and DBD+ cells (Supplementary Figure 6). However, in DBD+ cells, formation of two smaller TADs nested within the larger TAD is detected with an increased number of loops (Supplementary Figure 6B). DBD and DBD+ peaks cluster in similar locations across the hub. However, DBD+ binding at *MYEOV* and *CCND1* promoters were the only ones that effectively modulated chromatin at these microsatellites to promote gene expression (black tracks, padj < 0.05, FC >2). The *CCND1* promoter microsatellite has a density of 1 meaning that it only consists of GGAA repeats without any gaps, whereas *MYEOV* microsatellites densities were 0.44 and 0.28. We also found similar results at two more loci with EWSR1::FLI1-bound GGAA microsatellites: *NKX2-2* and *GSTM4* (Supplementary Figure 7 & 8) ***(Luo et al., 2009; Chong et al., 2018; Showpnil et al., 2022)***. These findings further demonstrate the preference of DBD-α4 helix for binding at dense GGAA repeats and the increased binding intensity seen for DBD without the DBD-α4 helix at shorter and less dense GGAA microsatellites. Taken together, these data provide examples of transcriptional hubs spanning a large genomic distance containing numerous genes and show how transcriptional output at these hubs can be regulated by EWSR1::FLI1 binding at GGAA microsatellites.

### The DBD-α4 helix is required for transcription and chromatin regulation in TTC-466 cells

In order to understand if these observations from A-673 cells are applicable in other models of Ewing sarcoma, we used our conventional KD/rescue system in TTC-466 cells, which express the EWSR1::ERG fusion. We depleted endogenous EWSR1::ERG with shRNA and rescued with DBD and DBD+ constructs and tested expression of rescue constructs, transcriptional changes and transformation in agar before performing Micro-C analysis (Supplementary Figure 9A-C). Importantly, the winged helix-turn-helix domain of wildtype FLI1 is highly homologous to ERG. From the first α1 helix through the flanking α4 helix of the ETS DBD, ERG varies from FLI1 in only two residues of α1 (alanines in FLI1 that are serines in ERG, ***Poon and Kim (2017)***). Our DBD and DBD+ constructs are thus >97% homologous with analogous constructs for ERG. These experiments largely recapitulated our results in A-673 cells (Supplementary Figure 1). Notably, PCA analysis of RNA-seq data showed that, as for A-673 cells, wtEF clustered away from the KD condition, with DBD+ clustering with wtEF and DBD clustering between KD and wtEF. (Supplementary Figure 9B and Supplementary Figure 1C). Taken together, these data suggest that the DBD-α4 helix is also required in transcription and transformation processes of Ewing sarcoma in TTC-466 cells.

Next, we sought to understand the 3D chromatin organization of TTC-466 cells expressing DBD and DBD+ proteins. First, we performed MDS analysis of the top 1000 interactions of Micro-C matrices at 500kb resolution (Supplementary Figure 10A). KD replicates clustered furthest away from DBD+ replicates. Additionally, DBD replicates fell between the KD and DBD+ replicates. We also assessed the contact frequency across the genome in combined replicates of KD, DBD and DBD+ cells at 5kb resolution (Supplementary Figure 10B). At short distance (up to around 10^5^ bp), cells expressing DBD have more contacts in comparison to KD and DBD+ cells, while DBD+ expressing cells make more frequent contacts than KD and DBD cells at longer distances (Supplementary Figure 10B). We further conducted differential interaction analysis using 500kb matrices to compare cells expressing DBD and DBD+ replicates to KD cells using the same cutoff values as above for A-673 cells. (Supplementary Figure 10C-D). First, we detected 42,891 and 2,598 interactions in DBD+ and DBD cells, respectively, using cutoffs of p-value < 0.2 and log_2_FC > 0.5. We then used more stringent criteria (adjusted p-value < 0.05, log_2_FC > 1.5) to identify DIRs. We found that DBD+ cells have 1,742 DIRs compared to KD cells (Supplementary Figure 10C) whereas DBD cells have 72 DIRs (Supplementary Figure 10D). Taken together, these results suggest that the DBD-α4 helix of FLI1 participates in restructuring 3D chromatin organization in TTC-466 cells.

We then repeated the TAD and loop level analysis in DBD and DBD+ expressing TTC-466 cells with respect to the KD condition. Surprisingly, we found more TADs and loops in DBD expressing cells (16,350 TADs; 1,646 loops) compared to DBD+ cells (7,099 TADs; 297 loops) with comparable patterns of overlap between DBD and DBD+ in both TAD and loop analysis (Supplementary Figures 11-12). Notably, despite a finding significantly more loops in the DBD condition, overlap analysis showed that the loops gained in DBD and DBD+ expression conditions are largely different while the loops lost in both conditions are similar (Supplementary Figure 12B-C). This pattern was also detected in A-673 cells (Supplementary 4A) suggesting that common mechanisms drive altered chromatin looping in both cell lines.

We next used CUT&Tag for H3K27ac and FLAG to assess the enhancer landscape and construct binding in DBD and DBD+ expressing TTC-466 cells (Supplementary Figure 13). Similar to A-673 cells (Figure 4A) and reflecting the transcriptional changes seen in RNA-seq (Supplementary Figure 9B), we found the H3K27ac signal for DBD samples clustered between DBD+ and KD conditions in TTC-466 cells (Supplementary Figure 13A). In contrast to A-673 cells, we found many more FLAG peaks in DBD+ expressing TTC-466 cells (61,665) compared to DBD expressing cells (21,000) with most DBD-bound loci commonly also bound by DBD+ (Supplementary Figure 13B). We further found that 55.29% of common peaks overlapped GGAA microsatellites, while 48.13% and 42.99% unique DBD and DBD+ peaks overlapping GGAA microsatellites, respectively (Supplementary Figure 13C). We then characterized the GGAA microsatellites uniquely bound by either DBD, DBD+, or both (Supplementary Figure 13D-13H). We found the common FLAG peaks were overlapped with the longest set of GGAA repeats (mean=37.79bp) compared to unique DBD peaks (mean=30.28bp) and DBD+ unique peaks (mean=22.74bp), and this led to common peaks having the longest and densest GGAA microsatellites in TTC-466 cells (Supplementary Figure 13D-H).

We then sampled *FCGRT*, *CCND1*, *NKX2-2*, and *GSTM4* in TTC-466 cells (Supplementary Figures 14-17, ***Gangwal et al. (2008b); Luo et al. (2009); Showpnil et al. (2022); Riggi et al. (2014); Chong et al. (2018)***). At these loci in TTC-466 cells, we found only few loops in *CCND1* and *GSTM4* regions (Supplementary Figures 14-17) and no TADs were detected, reflecting differences in our TAD and loop analysis between cell lines as outlined above and discussed below. However, the overall pattern of DBD and DBD+ binding was similar to that seen in A-673 cells, with DBD showing greater peak intensity at non-GGAA microsatellites and equal or lesser intensity at EWSR1::FLI1-bound GGAA microsatellites. Additionally, DBD+ expressing cells had greater H3K27ac peak intensity at these loci, also similar to that seen in A-673 cells and consistent with the rescue of transcriptional activity observed for DBD+ in both A-673 and TTC-466 cell lines. Thus, the genome-wide analyses and the Ewing-specific loci multiomic analysis using RNA-seq, Micro-C, H3K27ac CUT&Tag, and FLAG CUT&Tag indicate that the DBD-α4 helix is required for fusion-driven oncogenic gene regulation in Ewing sarcoma.

## Discussion

We found generally concordant results between cell lines, further supporting the critical role for the DBD-α4 helix in EWSR1::FLI1-mediated changes to transcription, the enhancer landscape, and 3D chromatin structure. There were a few differences between cell lines and this has many possible explanations, none of which are mutually exclusive. One possibility is the high level of heterogeneity found in genome regulation between Ewing sarcoma cell lines. Though there are several critical genes found to be regulated by EWSR1::FLI1 in many samples, very few genes are regulated the same way across all tested cell lines. As an example, high expression of *NKX2-2* can be used as a diagnostic marker in Ewing sarcoma and is upregulated by EWSR1::FLI1 binding at a distal GGAA microsatellite in many cell lines, including A-673 ***(Riggi et al., 2014; Showpnil et al., 2022)***. However, the chromatin architecture in TC71 cells has a unique TAD boundary insulating the *NKX2-2* gene from the EWSR1::FLI1-bound GGAA microsatellite and *NKX2-2* is instead highly expressed through other regulatory mechanisms ***(Showpnil et al., 2022)***. To address the cell line to cell line variability, we plotted RNA-seq of the replicates from both cell lines on a same PCA plot (Supplementary Figure 18A) and found that cell line specific gene expression comprised the majority of the variability, with each line clustering together along the first principal component. Notably, within each cell line similar clusters were found placing DBD+ near wtEF away from the KD condition along the second principal component, with DBD between these two clusters (Supplementary Figure 18A).

Another possible reason for differences, particularly in the TAD and loop analyses, is related to the sequencing depths in both cell lines. We plotted the sequencing depth of each replicate of Micro-C data in both cell lines (Supplementary 18B-C). In A-673 cells, the total reads were similar between biological replicates as well as different conditions: 900,000,000 reads (Supplementary 18B). For TTC-466 cells, there is more variability both between the replicates of the same condition as well as between conditions (Supplementary 18C). Specifically, the DBD+ replicates had total read of 450,000,000 (Supplementary Figure 18C, middle bars). This relatively low coverage of DBD+ replicates could contribute to lower numbers of TADs and loops detected in DBD+ TTC-466 cells compared to DBD TTC-466 cells. Lastly, another notable difference was found in the microsatellite binding preferences of different constructs in different cell lines. We found many more DBD+ FLAG peaks in TTC-466 cells. This may have captured many more binding events at smaller and less dense microsatellites. As it is difficult to analytically normalize for the possibility of cell-line specific factors that influence CUT&Tag, a more reductionist biochemical approach may be needed to fully resolve differences in affinities for GGAA microsatellites of varying characteristics between DBD and DBD+. Nonetheless, at specific loci regulated by EWSR1::FLI1 binding at GGAA repeats we found similar patterns of FLAG binding in A-673 and TTC-466. Specifically, DBD+ showed relatively higher peak intensity at long and dense GGAA microsatellites as compared to DBD. The opposite was true at the other shorter or non-GGAA microsatellite bound sites, with DBD showing a relatively higher peak intensity.

Detailed differential analysis of genomic binding patterns of DBD and DBD+ was not previously performed. In our previous study, we used CUT&RUN to identify DBD and DBD+ peaks and found 90% overlap ***(Boone et al., 2021)***. In this study, CUT&Tag showed that this overlap is only 60%. This provided us with an opportunity to discern meaningful difference in genomic localization and se-quence characteristics of bound loci across three groups: DBD unique, common, and DBD+ unique peaks. Importantly, the methods utilize different enzymes for the digestion step: Micrococcal nuclease (MNase) for CUT&RUN and Tn5 transposase for CUT&Tag. Thus, a few possible factors might explain the differences in peak detection between the two techniques. MNase enzyme has cleavage preferences at sites rich in adenylate, deoxyadenylate, and thymidylate ***(Cuatrecasas et al., 1967)***. It also shown to have a preference for open chromatin or nucleosome-free regions ***(Henikoff et al., 2011; Weiner et al., 2010)***. Because the specific effects of EWSR1::FLI1 binding on GGAA repeats in chromatin is unknown, these loci may be particularly susceptible to excess cleavage by MNase in a way that biases CUT&RUN results. In contrast, the Tn5 transposase is especially efficient at integrating adapters into open chromatin without chromatin degradation and may, therefore, be particularly efficient at capturing binding at GGAA repeat regions. Additionally, CUT&RUN generally has lower signals, higher background, and lower yields compared to CUT&Tag ***(Kaya-Okur et al., 2019)***. We thus attribute the difference in detection of FLAG peaks in our previous and current studies to the different enzymes and their ability to recognize and access repetitive elements such as GGAA repeats.

With the current results, we favor a model whereby the DBD-α4 helix stabilizes collective binding at high density GGAA microsatellites. Our findings underscore the specific characteristics of GGAA repeats bound by EWSR1::FLI1 to drive pathogenesis of Ewing sarcoma. At Ewing-specific gene loci such as *FCGRT* and *CCND1* (Figure 6 and Supplementary Figure 5), we found dense microsatellites. This finding was also recapitulated at the genome-wide level as seen in the preference of DBD+ unique peaks for denser microsatellites compared to common peaks (Figure 5G). The DBD+ unique peaks also show a preference of binding at microsatellites that are longer than the DBD unique peaks and shorter than the common peaks (Figure 5D) aligning with our previous findings that an optimal number of GGAA repeats is required for binding by EWSR1::FLI1 in Ewing sarcoma transformation ***(Johnson et al., 2017b)***.

Although our data demonstrates that the DBD-α4 helix is required in collective binding at GGAA repeats, we are still unable to decipher the exact mechanism of stabilization at such repeats with our current set of data. There are a few possible mechanisms. The DBD-α4 helix could be directly interacting with the DNA or the DNA binding domain of the adjacently bound EWSR1::FLI1 molecule. Another possibility is the DBD-α4 helix affects its function through interactions with other epigenetic regulators or transcription machinery proteins. Alternatively, intramolecular interaction with the EWSR1 domain may stabilize DNA binding and could further promote phase condensates. We favor the last mechanism since de-novo ability of EWSR1::FLI1 binding GGAA repeats depends on the presence of the EWSR1 domain ***(Johnson et al., 2017a)***. Further directed studies are needed to address these possible mechanisms.

The multi-omics approach utilized in this study provided us with an opportunity to characterize transcription hubs driven by EWSR1::FLI1 genome-wide. We have shown the clustering of EWSR1::FLI1 at GGAA microsatellites underlies the formation of local 3D features such as TADs and chromatin loops. If we use TADs as proxies for hubs, we detected thousands more of these hubs in DBD+ cells compared to DBD cells, highlighting the importance of DBD-α4 helix in binding at dense GGAA repeats and formation of hubs across the genome. We also observed thousands of loops at unique microsatellites for both DBD and DBD+ cells adding detail to the architecture of hubs, often represented as flower-shaped structures of many loops ***(Zhu et al., 2021)***. We also probed another aspect of hubs with our H3K27ac CUT&Tag data: the presence of enhancers and super-enhancers. We showed that the DBD-α4 helix promotes more active marks genome-wide compared to DBD leading to formation of enhancers and super-enhancer. Finally, we presented select loci as specific examples of productive transcription hubs driven by EWSR1::FLI1 binding at dense GGAA microsatellites. Our study thus suggests a surprising role for FLI1 DBD in the process of hub formation which is usually attributed to the EWSR1 low complexity domain.

Because EWSR1::FLI1 is the sole driver mutation in Ewing sarcoma tumors, it is an attractive therapeutic target. However, many approaches to target EWSR1::FLI1 have been hampered by common issues that arise with targeting transcription factors, such as its location in the nucleus, its abundance in cells, and its lack of an enzymatic pocket to design a small molecule tailored to target it ***(Knott et al., 2019)***. We propose the DBD-α4 helix is a promising therapeutic target as it doesn’t directly bind the major groove of the DNA. Additionally, because the DBD-α4 is a helix, it provides a structured region to design small molecules to disrupt its interaction unlike the EWSR1 low complexity domain. Moreover, its importance in maintaining effective binding at GGAA microsatellites offers an opportunity to target EWSR1::FLI1 at the most mechanistically important sites in pathogenesis of Ewing sarcoma. Further studies are needed to clarify the mechanism by which the DBD-α4 helix promotes effective binding at GGAA microsatellites and regulates transcription hub formation.

## Materials and Methods

### Constructs and retroviruses

Mammalian expression constructs used include: Retroviral vectors encoding shRNA for luciferase-RNAi, EWSR1::FLI1-RNAi and EWSR1::ERG-RNAi as well as cDNA-containing vectors encoding 3xFLAG-EF, 3xFLAG-EF DBD, 3xFLAG-EF DBD+ ***(Boone et al., 2021)***. The EWSR1::FLI1 DBD, and EWSR1::FLI1 DBD+ were ordered as gene blocks (Integrated DNA Technologies) and cloned into the pMSCV-hygro plasmid between BamHI and AgeI restriction sites.

### Cell culture methods

HEK293-EBNA cells were grown at 37^◦^C, 5% CO2 in Dulbecco’s modified Eagle’s medium (DMEM, Corning Cellgro 10-013-CV), with 10% heat-inactivated fetal bovine serum (Gibco 16000-044), 1% penicillin/streptomycin/glutamine (P/S/Q, Gibco 10378-016), and 0.3 mg/Ml Geneticin (Gibco 10131-027). A-673 was obtained from American Type Culture Collection (ATCC, Manassas, VA). These cells were grown at 37C, 5% CO2 in DMEM with 10% fetal bovine serum, 1% P/S/Q, and 1% sodium pyruvate (Gibco 11360-070). TTC-466 cells were obtained from Timothy Triche, MD, PhD (CHLA) and were grown at 37C, 5% CO2 in RPMI with 10% fetal bovine serum, and 1% P/S/Q. For knockdown of endogenous EWSR1::FLI1 in A-673 and EWSR1::ERG in TTC-466, cells were infected with RNAi virus and subsequently infected with the cDNA-containing virus to rescue the cells. After 48 hours, cells were selected with puromycin (100 µg/ul) and hygromycin (150 µg/ul); and allowed to grow for 7-8 days prior to collection for downstream analysis. Cells were tested regularly for Mycoplasma using the PCR based Universal Mycoplasma Detection Kit (ATCC, 30-1012K). Cell line identities were confirmed by short tandem repeat (STR) profiling (Genetica LabCorp, USA), last performed in February 2022.

### Transfection, virus production and transduction

For the generation of retroviruses, HEK293-EBNA cells were co-transfected with retroviral expression plasmids, vesicular stomatitis virus G glycoprotein (VSV-G) and gag/pol packaging plasmids. Briefly, 2.5 x 106 HEK293-EBNA cells were seeded in a 10 cm tissue culture dish the day before transfection, resulting in 60-70% confluency the day of transfection. 10 µg of each plasmid (gag-pol, vsv-g, and transfer plasmid) were combined with 2 ml Opti-MEM I Reduced Serum Medium (Gibco 31985070) and 90 µl MirusBio TransIT-LT1 Transfection Reagent (Mirus MIR2306) and incubated at room temperature for 20 minutes. The transfection mix was then added drop wise to the cells in 3 ml culture medium. Virus-containing supernatant was collected every 4 hours on day 2 (48 hours) and 3 (72 hours) post transfection, pooled, filtered and stored at –80°C. Ewing Sarcoma cells were transduced with viral supernatants using polybrene (8 µg/ml), followed by selection with appropriate antibiotics at 48 hours post infection. In the case of knockdown/rescues, cells were selected with 0.5-2 µg/ml Puromycin and 50-150 µg/ml Hygromycin B (Thermo 10687010) for 7-10 days.

### Immunodetection

Whole-cell protein extraction was completed using Pierce™ RIPA buffer (ThermoFisher 88901) supplemented with Protease Inhibitor Cocktail (Sigma P8340-5ML) on ice for 30 minutes. For nuclear extracts cell pellets were resuspended in LB1 buffer (50 mM Tris-HCl pH 7.5, 20 mM NaCl, 1 mM EDTA, 0.5% NP-40, 0.25%, Triton-X 100, 10% Glycerol, 1 mM DTT, Protease Inhibitor Cocktail) and incubated on a nutator for 10 minutes at 4C. After a 5 minute spin at 400xg and 4C, the pellets were washed with LB2 buffer (10 mM Tris-HCl pH 7.5, 20 mM NaCl, 1 mM EDTA, 0.5 mM EGTA, 1 mM DTT, Protease Inhibitor Cocktail) and centrifuged again for 5 minutes at 400xg and 4C. The nuclear pellets were resuspended in a small volume of RIPA buffer, incubated on ice for 30 minutes, followed by a 30 minute centrifugation step at maximum speed and 4C.

Protein concentration was determined using Pierce™ BCA™ Protein Assay Kit (Thermo Scientific 23225). 15-35 µg of protein samples were ran on precast 4-15% gradient gels (Bio Rad) and transferred to nitrocellulose membranes (Invitrogen). 4-15% Mini-PROTEAN® TGX™ precast protein gel (Bio-Rad 4561084) and resolved at 120 V. Proteins were transferred to a nitrocellulose membrane (Thermo IB23002) using the iBlot™ 2 Gel Transfer Device (Thermo IB21001). The membrane was blocked with Odyssey® PBS Blocking Buffer (Li-Cor 927-40003) for 1 hour at room temperature. Immunoblotting was performed overnight at 4C using the following primary antibodies: anti-FLI1 rabbit (Abcam ab15289, 1:1000); monoclonal anti-FLAG M2 mouse (Sigma F1804-200UG, 1:1000); anti--Tubulin [DM1A] mouse (Abcam ab7291, 1:2000), and anti-Lamin B1 [EPR8985(B)] rabbit (Abcam ab133741, 1:1000). The membrane was washed with TBS/0.1% Tween 20 (TBS-T) and incubated with IRDye secondary antibodies (IRDye® 680LT donkey anti-rabbit IgG, IRDye® 800CW goat anti-rabbit IgG (H + L), IRDye® 800CW goat anti-mouse IgG1 specific secondary antibody, Li-Cor 926-68023, 926-32211, 926-32350, 1:2000) for 1 hour at room temperature. After a final wash step with TBS-T the membrane was imaged using the Li-Cor Odyssey CLx Infrared Imaging System. ImageJ software was used to perform densitometry analysis.

### Cyclohexamide chase assay

A6327 cells were treated with 50 µg/mL cycloheximide (Cayman 14126) to assess protein stability. A 50 mg/mL CHX stock solution was freshly prepared in dimethyl sulfoxide (Thermo Fisher D12345), and an equivalent volume of DMSO was added to generate time zero samples. After 0, 2, 4, 6, 18 and 24 hours, cells were harvested, then washed once in ice-cold Hank’s Balanced Salt Solution (Gibco 14025134) and pelleted at 400xg for 5min at 4C. The cell pellets were lysed on ice in NP-40 buffer (50 mM Tris-HCl, pH 8.0, 150 mM NaCl, 1% NP-40) supplemented with phenylmethylsulfonyl fluoride (Thermo Fisher 50-165-6975) for 30min. The total lysates were clarified by centrifugation at 12,000 × g for 15 minutes at 4C. Protein concentration was determined using the Pierce™ BCA™ Protein Assay Kit (Thermo Scientific 23225). Samples were stored at 80C until immunoblotting. Band intensities were quantified using ImageJ (NIH), and protein half-lives were calculated from signal decay curves fitted in GraphPad Prism (v10.0, GraphPad Software).

### Soft agar assays

The anchorage-independent growth capacity of Ewing sarcoma cells was assessed using soft agar assays. Cells were seeded at a density of 7500 cells in 6-cm3 tissue culture dish in duplicate in 0.8% SeaPlaque GTG agarose (Lonza 50111) mixed with Iscove’s Modified Dulbecco’s medium (Gibco 12200-036) containing 20% FBS, penicillin/streptomycin/glutamine and puromycin/hygromycin. Agars were imaged at least 14 days after seeding and colony counts were quantified using ImageJ software (V1.51).

### RNA-sequencing experiments, data processing, and analysis

RNA-sequencing was performed on 3 biological replicates of KD, DBD and DBD+. Total RNA were extracted using RNeasy Extraction Kit and submitted to the Nationwide Children’s Hospital Institute for Genomic Medicine for RNA quality measurement, library preparation, and sequencing. Briefly, cDNA libraries were prepared from total RNA with TruSeq Stranded mRNA Kit (Illumina 20020594) and sequenced on Illumina NovaSeq SP to generate 150-bp paired-end reads. We used in-house RNA-sequencing pipeline to process and analyze the data. Low-quality reads (q < 10) and adapter sequences were trimmed to align to hg19 genome using STAR ***(Dobin et al., 2013)***. After alignment, the reads were counted and differential analysis performed using DESeq2 ***(Love et al., 2014)***.

### CUT&Tag experiments

CUT&Tag was performed as described in ***Kaya-Okur et al. (2019)*** with slight modifications. 250,000 cells per CUT&Tag condition were bound to BioMag® Plus Concanavalin A-coated magnetic beads (Bangs Laboratories, BP531) and incubated with the primary antibody (anti-FLAG M2 mouse, Sigma F1804-200UG, 1:100) overnight at 4 °C and secondary antibody (rabbit anti-mouse, Abcam ab46540, 1:100) for 1 hour at room temperature.

Adapter-loaded protein A-Tn5 fusion protein was added at a dilution of 1:250 and incubated for 1 hour at room temperature. To activate the Tn5, tagmentation buffer containing MgCl2 was added and samples were incubated for 1 hour at 37°C. Reactions were stopped by addition of EDTA and DNA was solubilized with SDS and Proteinase K for 1 hour at 50°C. Total DNA was purified using Phenol/Chloroform extraction followed by Ethanol precipitation. CUT&Tag libraries were prepared with NEBNext HiFi 2x PCR Master Mix (NEB M0541S) and indexed primers ***(Buenrostro et al., 2015)*** using a combined Annealing/Extension step at 63°C for 10 seconds and 15 cycles followed by a 1.1X post-amplification AMPure XP (Beckman Coulter, A63880) bead cleanup. The fragment size distributions and concentrations of the final libraries were determined using the High Sensitivity D1000 Screen tape assay and reagents (Agilent, 5067-5584 and 5067-5585) on the Agilent 2200 TapeStation System. Libraries were pooled and sequenced (2 x 150 bp paired end) on the Illumina NovaSeq S1-Xp system (Nationwide Children’s Hospital Institute for Genomic Medicine).

### Micro-C experiments

Micro-C kits (Catalog 21006) purchased from Dovetail Genomics were used to prepare Micro-C libraries. For each condition, multiple aliquots of 1*x*10^6^ cells were harvested and frozen at –80°C for at least 30 minutes. Cells were thawed in room temperature and resuspended in first PBS containing 0.3M DSG then in 37% formaldehyde to crosslink DNA. Cells were then digested with various amount of MNase to achieve the digestion profile of 40%-70% mononucleosome peak observed on TapeStation D5000 HS Screen Tape. Conditions that are to be analyzed comparatively were digested to a similar range of mononucleosome peaks (50%-70%). Once desired digestion profiles achieved, the cells were lysed and the chromatin was captured with beads to perform proximity ligation. Libraries were prepared per the protocol of Micro-C kit and each library was indexed with unique primer pairs from IDT (10009816 and 10010147). Micro-C libraries were then shallow sequenced at 7 to 8 million (2 x 150bp) read pairs on Illumina NovaSeq6000 and the QC analysis pipeline provided from Dovetail Genomics were used to assess the quality of each library. Libraries that passed the QC step was then sequenced up to 300 million read pairs on NovaSeq6000.

### CUT&Tag data processing and analysis

CUT&Tag experiments were carried out for 2 biological replicates of CTCF and H3K27ac and 3 biological replicates for FLAG tagged DBD and DBD+ constructs. An in-house pipeline was used to analyze CUT&Tag data ***(Boone et al., 2021; Showpnil et al., 2022)***. Quality control on raw sequencing reads were performed with FastQC (v0.11.4) ***(Andrews, 2010)***. Adapter sequences and/or low quality reads were trimmed using trim_galore (0.4.4_dev) ***(Krueger, 2015)***. Reads were aligned to human (hg19) and spike-in Escherichia coli (Escherichia_coli_K_12_DH10B NCBI 2008-03-17) genomes using Bowtie2 (v2.3.4.3)11,12 with the following options ‘–no-unal –no-mixed –no-discordant –dovetail –phred33 –q –I 10 –X 700’. ‘–very-sensitive’ option was added when aligning to spike-in genome ***(Langmead and Salzberg, 2012)***. SamTools (v1.9) was used to convert sam to bam with ‘-bq 10’ option ***(Li and Durbin, 2009)***. CUT&Tag reads were spike-in normalized using DESeq2’s median ratio method to eliminate bias across different samples, minimize the effect of few outliers and appropriately account for global occupancy changes ***(Anders and Huber, 2010)***. Spike-in normalized tracks were generated and averaged across biological replicates using deepTools ***(Ramirez et al., 2016)***. Peaks were called with spike-in normalization and their corresponding IgG as controls accounting for variation between the biological replicates using MACS2 (v 2.2.7.1) ***(Zhang et al., 2008)***, DiffBind (v2.14.0) ***(Ross-Innes et al., 2012; Stark R., 2011)*** and DESeq2 (v1.26.0) ***(Love et al., 2014)***. All duplicate reads were kept in the analysis. Irreproducible Discovery Rate (IDR) (v 2.0.3) ***(Li et al., 2011)*** was used to identify reproducible and consistent peaks across replicates. To ensure high quality peaks that are most likely to represent biological signals, the final peak lists were generated with following default thresholds: FDR < 0.05, log2 Fold-Change > 8, mean normalized counts of signal > 80, and IDR < 0.01.

### Micro-C data processing and analysis

Micro-C libraries of 2 biological replicates of KD, DBD and DBD+ were prepared. Sequenced libraries were processed per instructions of Dovetail Genomics. Briefly, fastq files were aligned to hg19 reference genome using BWA-MEM algorithm with options –5SP to map mates independently ***(Li and Durbin, 2010)***. Next, parse module from pairtools (**?**) was used to find ligation junctions in Micro-C libraries with options min-mapq 40 (alignment with mapq < 40 will be marked as multi) and max-inter-align-gap 30 (if the gap is 30 or smaller, ignore the map, if the gap is >30, mark as “null” alignment). The parsed pair is then sorted using pairtools sort and PCR duplicates were removed with pairtools dedup. The pairtools split command was used to split the final .pairsam into .bam (then sorted with samtools sort) and .pairs files. Using Juicer Tools, .pairs files were converted into HiC contact matrices ***(Durand et al., 2016b)***. HiC matrices were then converted to mcool matrices using hic2cool ***(Durand et al., 2016a)***.

For MDS plot of individual replicates (Figure 1B), 500kb resolution cool matrices were converted gi interaction object using hicConvertFormat from HiCExplorer (v.3.7.2) ***(Wolff et al., 2020)***. Then using diffHiC package ***(Lun and Smyth, 2015)***, bins with low average abundance and low absolute counts were filtered out. Filtered reads were then scaled using library size and bin pairs that were on the diagonal line were also removed from analysis. Joint normalization of all replicates were carried out with diffHiC. Specifically, normOffsets function was used to remove trended biases with loess normalization and then a new set of log2-transformed counts adjusted by the negative binomial offset were computed. Batch effect was removed using removeBatchEffect from limma package ***(Ritchie et al., 2015)*** before plotting the top 1000 interactions. For distance-decay plot (Figre 1C), individual replicates were combined. First, the combined matrices were normalized using hicNormalize function from HiCExplorer to scale the libraries to the smallest library ***(Wolff et al., 2020)***. Scaled libraries were then plotted for diagnostic plots to determine the thresholds to use in hicCorrectMatrix function for ICE normalization. ICE-corrected 5kb matrices were then plotted with hicPlotDistVsCounts. For differentially interacting region analysis (Figure 1D-F), multiHiCcompare package was used ***(Stansfield et al., 2019)***. Briefly, individual replicates of KD, DBD and DBD+ were loess normalized and pairwise comparison of DBD+ to KD and DBD to KD was done using QLF (quasi-likelihood) method with batch effect correction. Volcano plots of differentially interacting regions with padj < 0.05 and fold-change > 1.5 plotted for each comparison (DBD+ and DBD vs KD).

For TAD analysis, individual replicates were combined and then scaled using the smallest library size with hicNormalize from HiCExplorer (v.3.7.2) ***(Wolff et al., 2020)***. Then matrices at 10kb, 25kb, 50kb and 100kb resolutions were ICE-corrected with the thresholds determined from diagnostic plots. First, using hicFindTADs function, we called TADs at the previously mentioned 4 resolutions for DBD and DBD+ matrices. Then, hicDifferentialTAD used to compute differential TADs by comparing the precomputed DBD and DBD+ TAD regions with the same regions of KD matrix. Differential TADs from each resolution were combined using hicMergeDomains with default –value of 5000 to account for duplicated TADs. For CUT&Tag peak annotation, peaks (FDR < 0.05, log2 Fold-Change > 8, mean normalized counts of signal > 80, and IDR < 0.01) were overlapped with findOverlaps functions from GenomicRanges package ***(Lawrence et al., 2013)***.

For loop calling, combined replicates at 1kb resolution matrices that were scaled and normalized in same manner for TAD analysis were used with Mustache (v.1.2.0) ***(Roayaei Ardakany et al., 2020)***. Mustache uses scale-space theory in computer vision to detect chromatin loops. For differential loops compared to KD matrix, diff_mustache.py was used to detect loops that were gained in DBD and DBD+ compared to KD and loops that were lost in DBD and DBD+ compared to KD matrix.

### Statistical analysis

When comparing means of two groups, two-sided Student’s t test was used. When comparing more than two groups, ANOVA test was used with Tukey honest significant differences test. * P value < 0.05, ** P value < 0.01, and *** P value < 0.001

## Supplementary Data

**Supplementary Figure 1:**
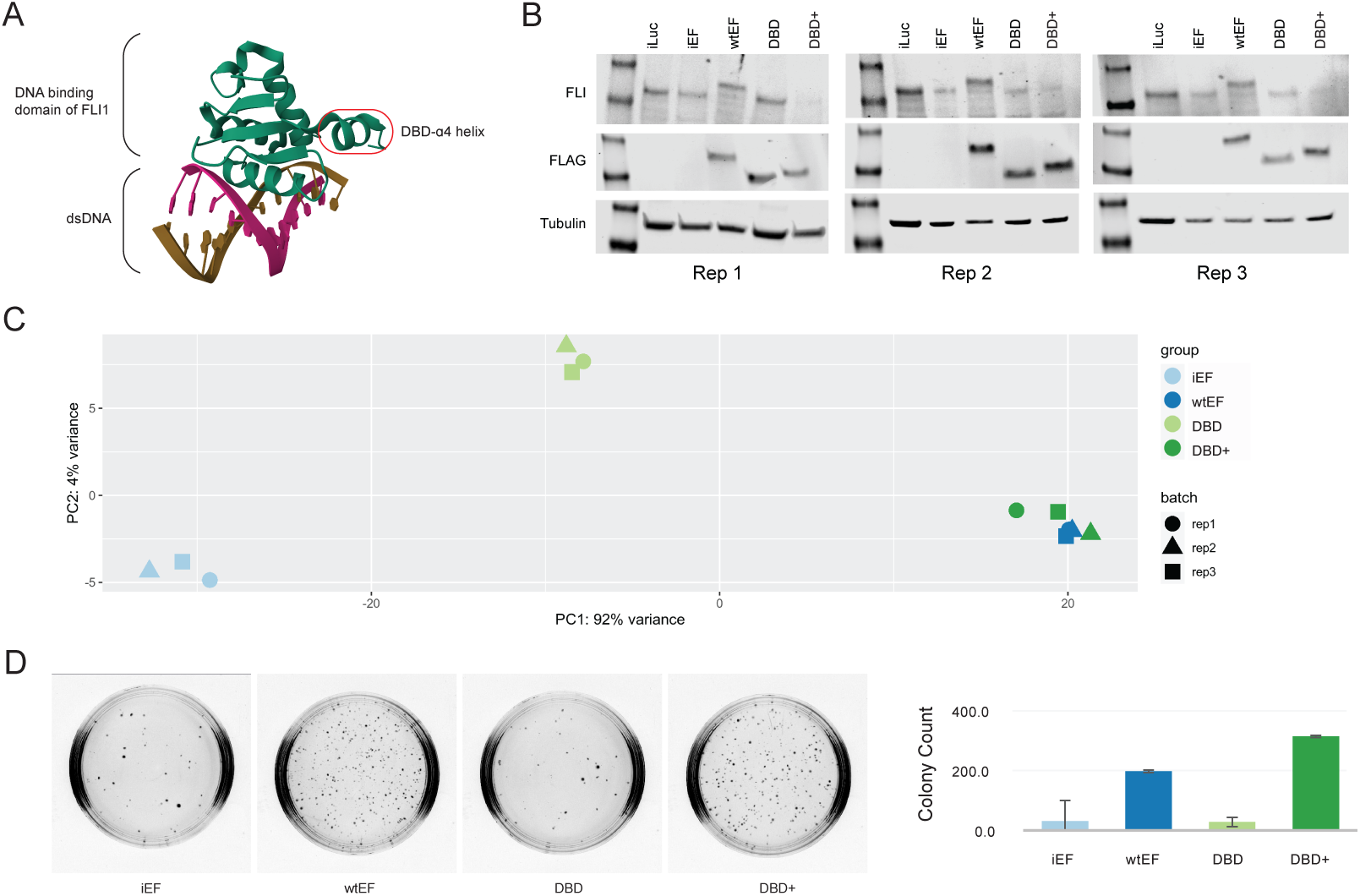
A. The DBD-α4 helix of FLI1 depicted on dsDNA (PDB) B. Knockdown of endogenous EWSR1::FLI1 detected with FLI1 ab and rescue of wtEF, DBD, and DBD+ detected with FLAG ab. C. A PCA plot of RNA-seq experiments in A-673 cells. D. Representative image of soft agar colony plates and quantification of three biological replicates.

**Supplementary Figure 2.**
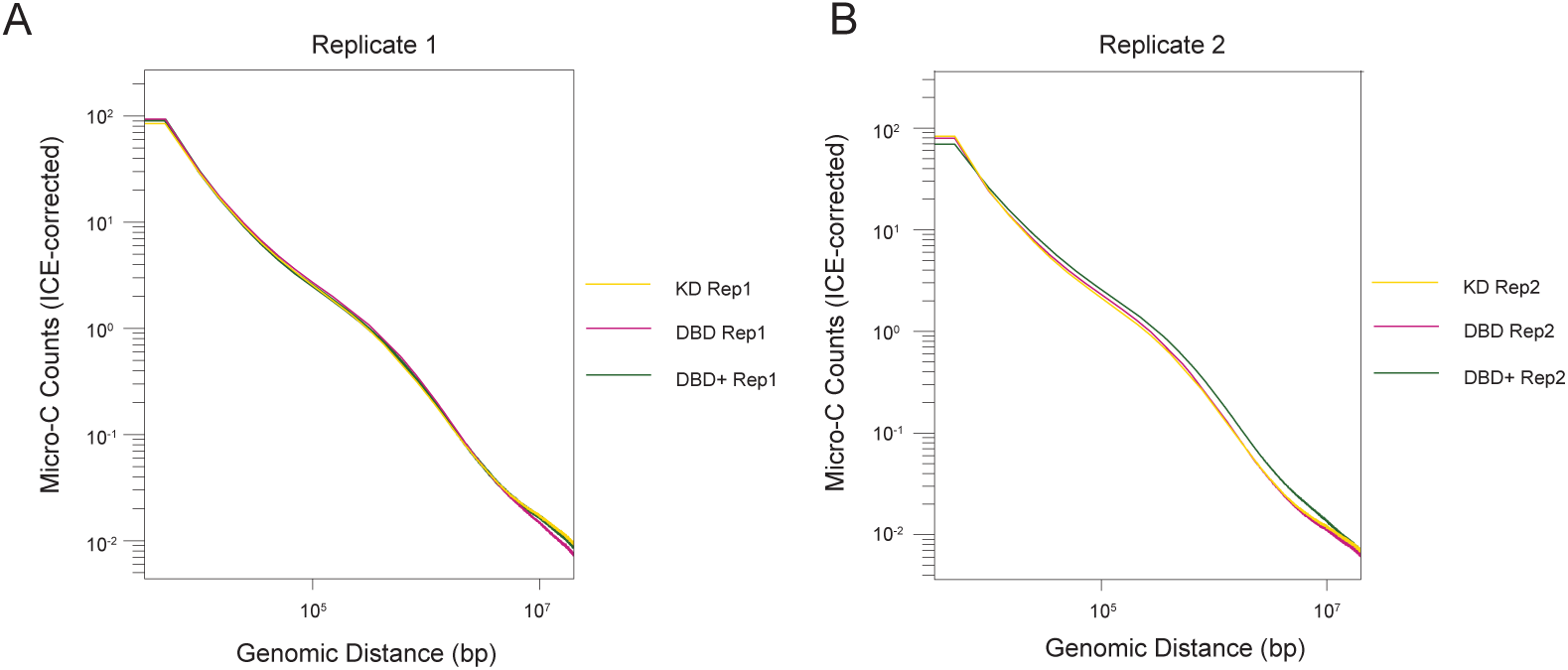
A. Genome-wide interaction frequency (ICE-corrected Micro-C counts) over genomic distance (bp) at 5kb resolution for Replicate 1 of A-673. B. Genome-wide interaction frequency (ICE-corrected Micro-C counts) over genomic distance (bp) at 5kb resolution for Replicate 2 of A-673.

**Supplementary Figure 3.**
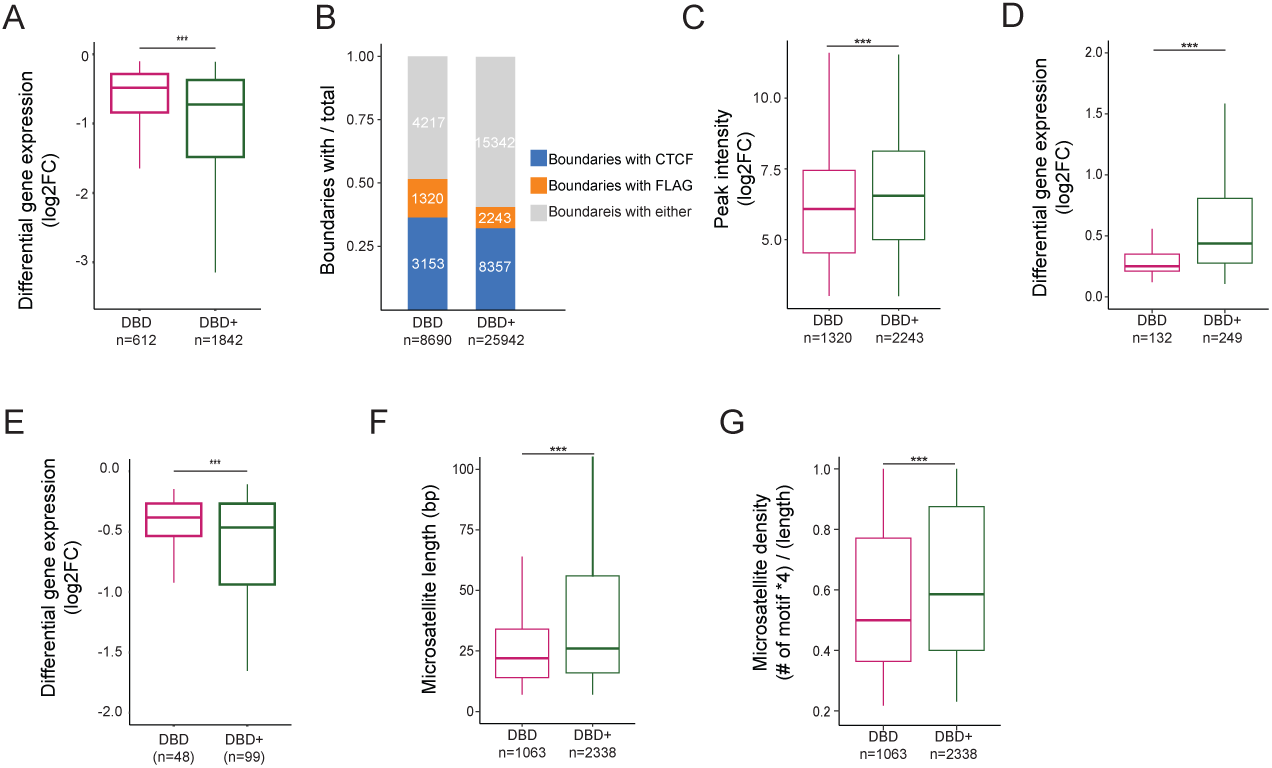
A. Expression level of significant genes overlapped with unique TADs in DBD (mean=-0.64) and DBD+ (mean=-1.16) bound by FLAG at GGAA microsatellites. B. Proportion of TAD boundaries bound by FLAG, CTCF or neither. C-G. Comparison of DBD and DBD+ unique TAD boundaries. C. Binding intensity of unique FLAG peaks (FDR < 0.05, FC > 8, counts > 80, IDR < 0.01) at boundaries of DBD and DBD+ unique TADs. D. Expression level of significantly upregulated genes ovarlapped with boundaries of unique TADs in DBD and DBD+ bound by FLAG at GGAA microsatellites. E. Expression level of significantly downregulated genes ovarlapped with boundaries of unique TADs in DBD and DBD+ bound by FLAG at GGAA microsatellites. F. Length of microsatellites bound by unique FLAG peaks at the boundaries of DBD and DBD+ conditions in bp. G. Percent of GGAA motif in the microsatellites calculated as (# of motif x 4)/(length of microsatellites) at the boundaries of DBD and DBD+ unique TADs bound by unique FLAG peaks. Boxplots depict the minimum, first quartile, median, third quartile, and maximum. *** P value < 0.001

**Supplementary Figure 4.**
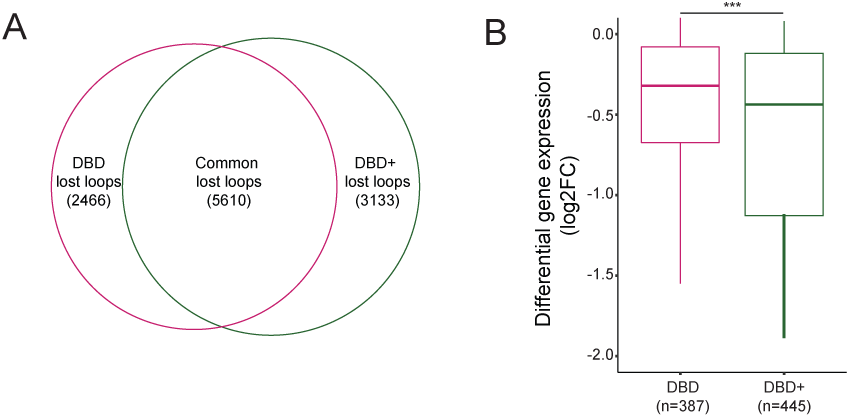
A. Venn diagram of overlap between DBD and DBD+ uniquely lost loops (compared to KD). B. Expression level of downregulated genes overlapped with uniquely gained loop anchors of DBD and DBD+. Means = –0.68, –1.25, 0.35, –1.08 * P value < 0.05, *** P value < 0.001. Boxplots depict the minimum, first quartile, median, third quartile, and maximum.

**Supplementary Figure 5.**
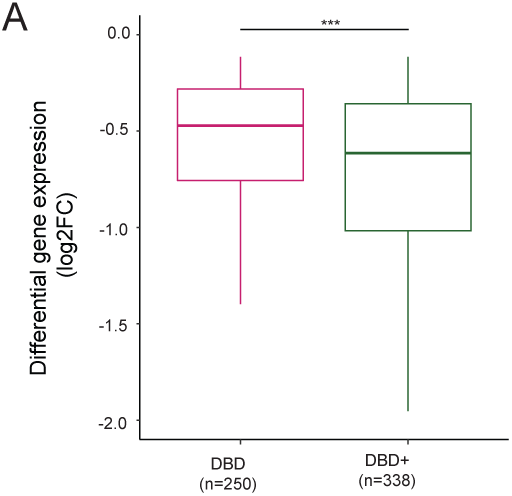
A. Expression level of downregulated genes at DBD and DBD+ super enhancers. *** P value < 0.001 Boxplots depict the minimum, first quartile, median, third quartile, and maximum.

**Supplementary Figure 6.**
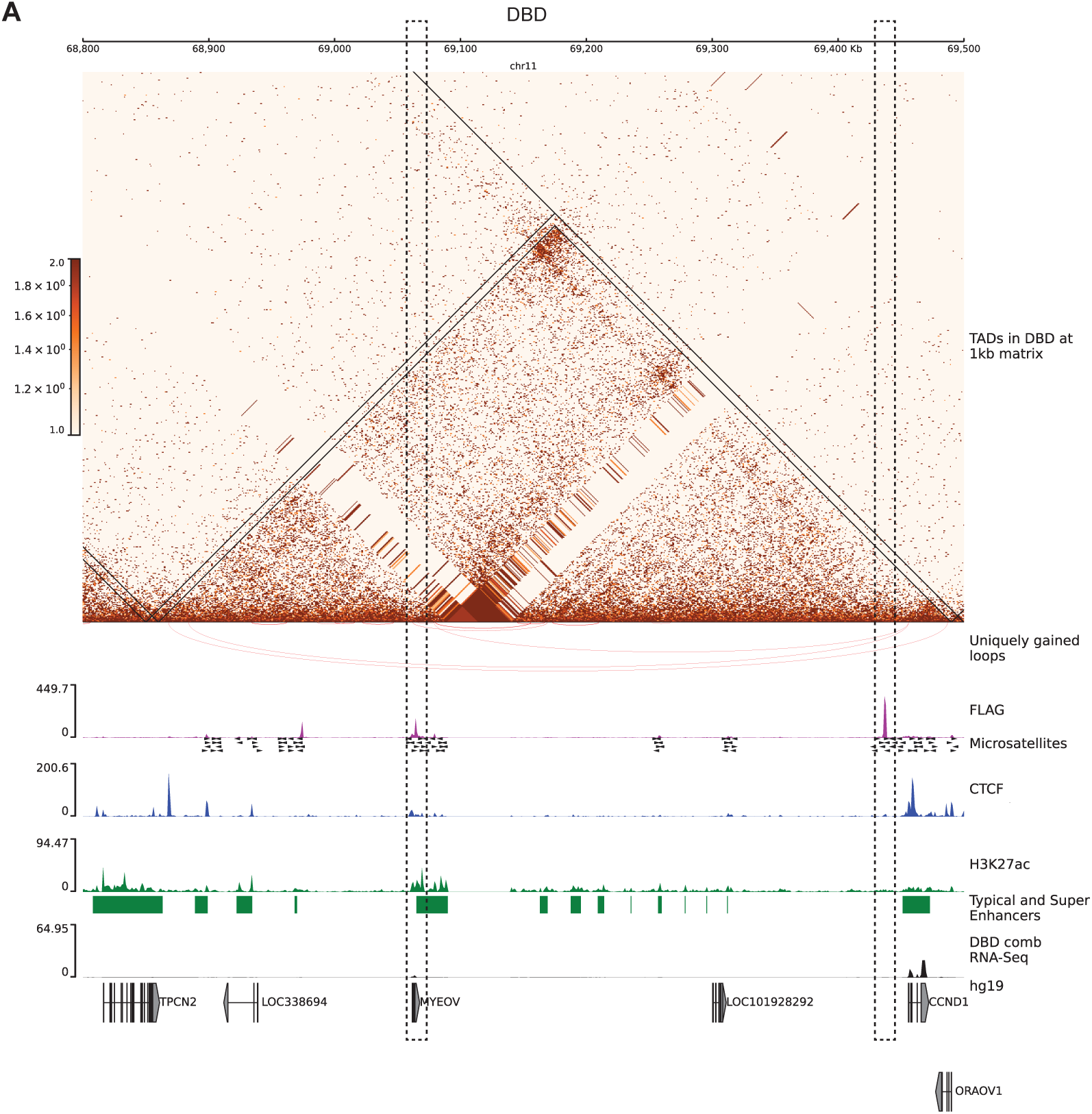
A. *CCND1* hub in 700kb region on chr 11 in A-673 DBD cells. TADs are depicted on 1kb matrices (DBD/KD). Uniquely gained loops are shown as red inverted arcs. FLAG CUT&Tag bigwig tracks depicted in magenta. GGAA microsatellites in hg19. CTCF CUT&Tag track is in blue middle row. H3K27ac tracks are in green. Enhancers and super-enhancers are shown as green bars. Gene expression is in black tracks.

**Supplementary Figure 6.**
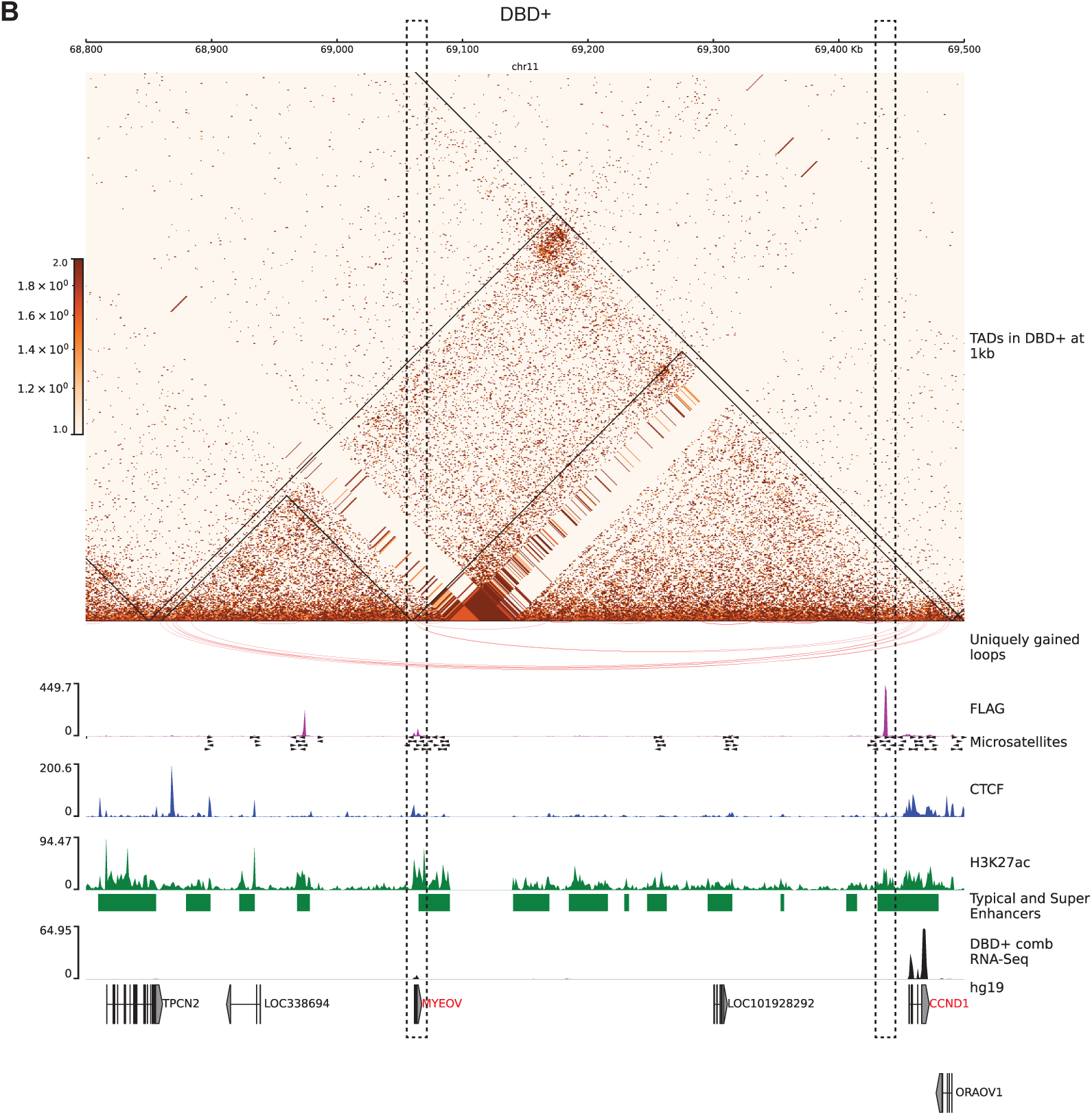
B. *CCND1* hub in 700kb region on chr 11 in A-673 DBD+ cells. TADs are depicted on 1kb matrices (DBD+/KD). Uniquely gained loops are shown as red inverted arcs. FLAG CUT&Tag bigwig tracks depicted in magenta. GGAA microsatellites in hg19. CTCF CUT&Tag track is in blue middle row. H3K27ac tracks are in green. Enhancers and super-enhancers are shown as green bars. Gene expression is in black tracks.

**Supplementary Figure 7.**
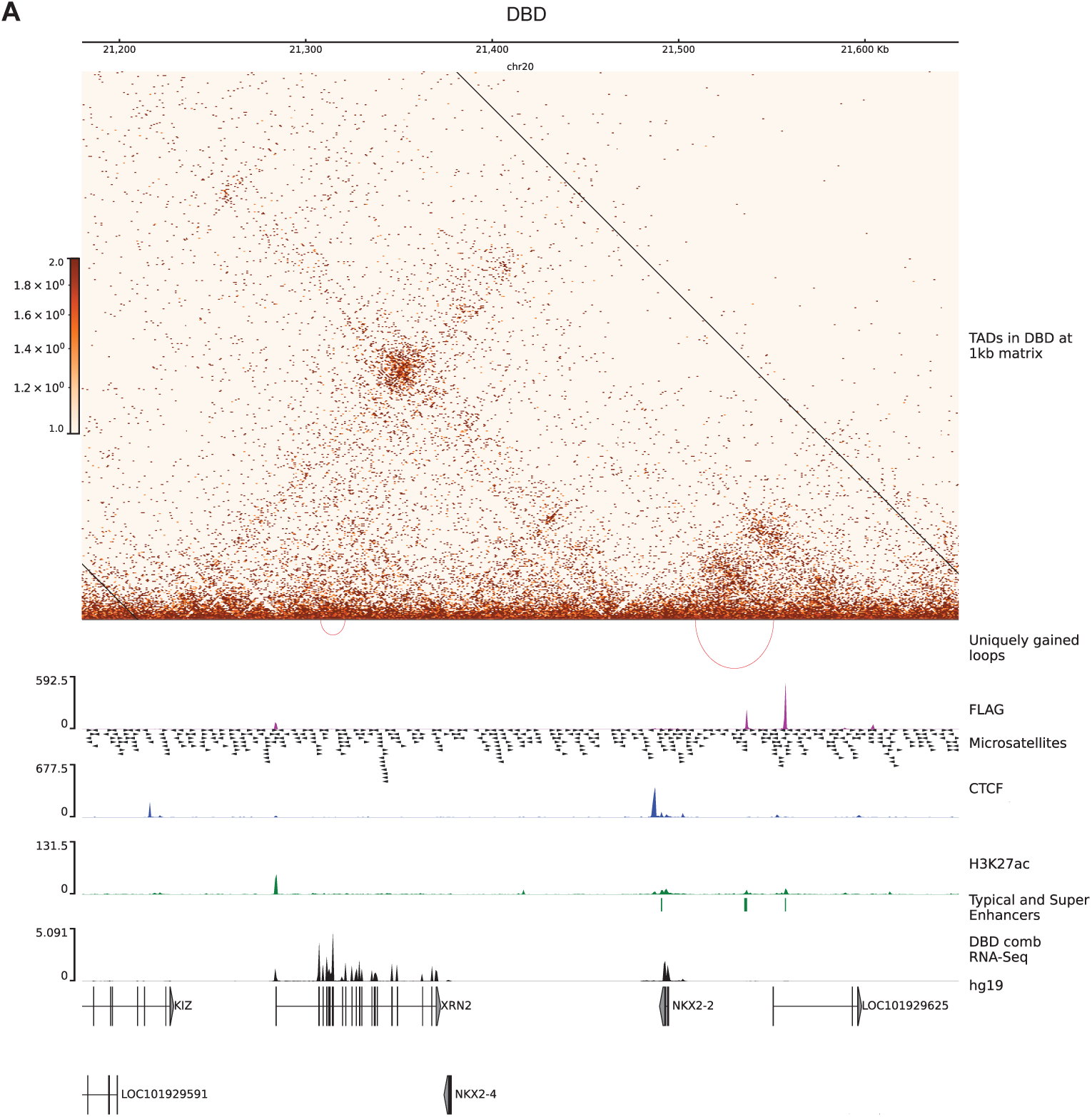
A. *NKX2-2* hub on chr 20 in A-673 DBD cells. TADs are depicted on 1kb matrices (DBD/KD). Uniquely gained loops are shown as red inverted arcs. FLAG CUT&Tag bigwig tracks depicted in magenta. GGAA microsatellites in hg19. CTCF CUT&Tag track is in blue middle row. H3K27ac tracks are in green. Enhancers and super-enhancers are shown as green bars. Gene expression is in black tracks.

**Supplementary Figure 7.**
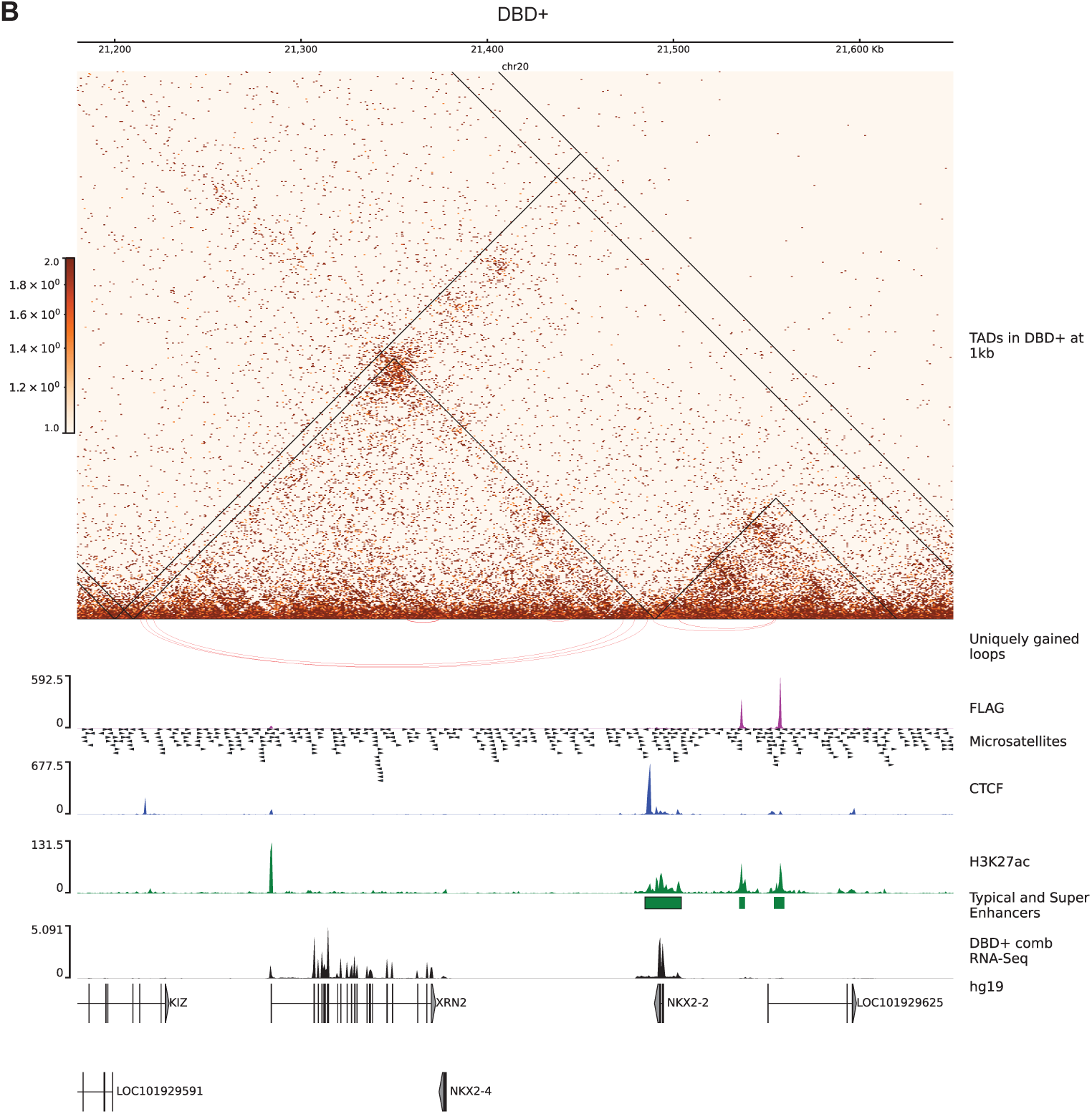
B. *NKX2-2* hub on chr 20 in A-673 DBD+ cells. TADs are depicted on 1kb matrices (DBD+/KD). Uniquely gained loops are shown as red inverted arcs. FLAG CUT&Tag bigwig tracks depicted in magenta. GGAA microsatellites in hg19. CTCF CUT&Tag track is in blue middle row. H3K27ac tracks are in green. Enhancers and super-enhancers are shown as green bars. Gene expression is in black tracks.

**Supplementary Figure 8.**
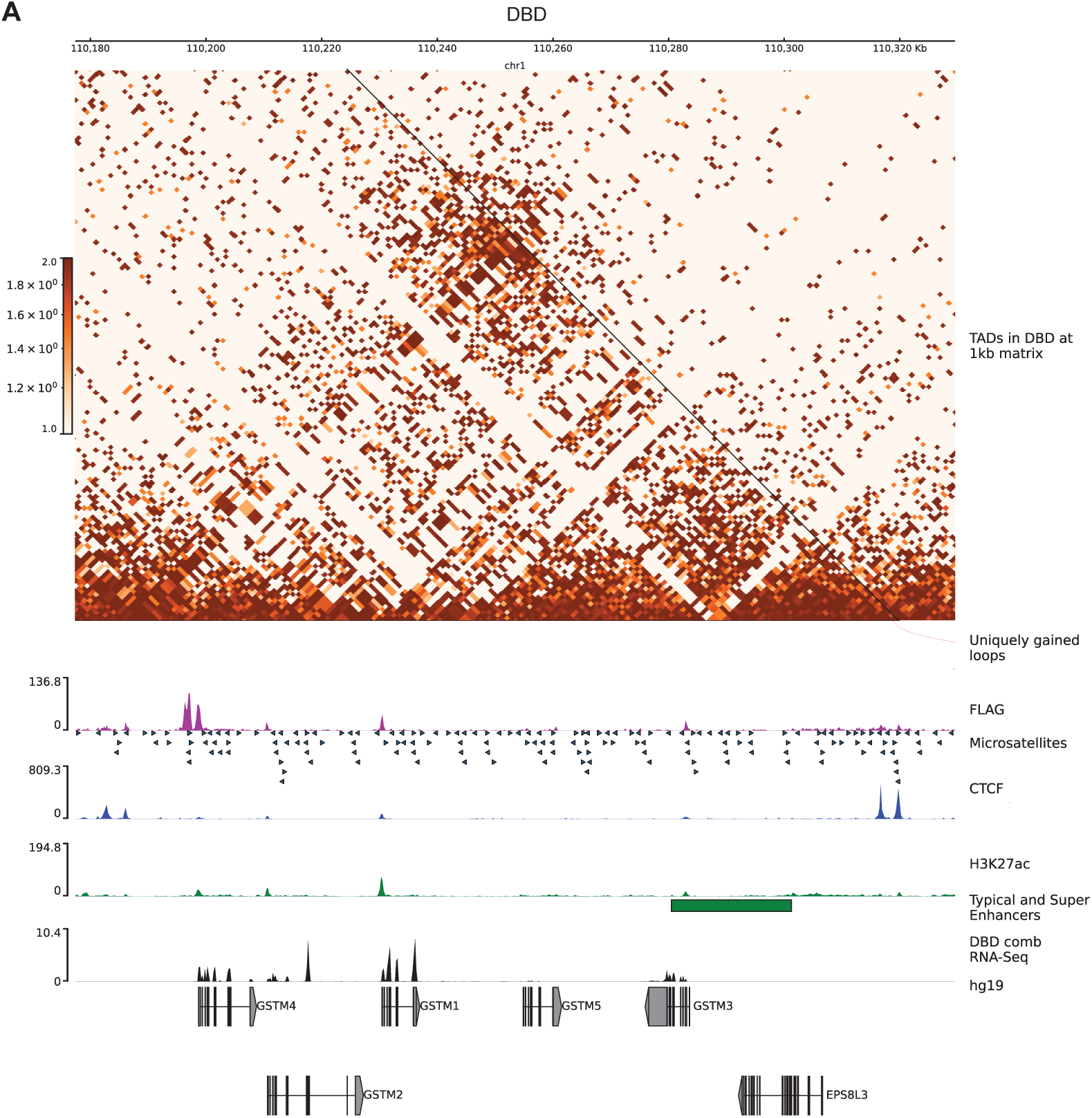
A. *GSTM4* hub chr 1 in A-673 DBD cells. TADs are depicted on 1kb matrices (DBD/KD). Uniquely gained loops are shown as red inverted arcs. FLAG CUT&Tag bigwig tracks depicted in magenta. GGAA microsatellites in hg19. CTCF CUT&Tag track is in blue middle row. H3K27ac tracks are in green. Enhancers and super-enhancers are shown as green bars. Gene expression is in black tracks.

**Supplementary Figure 8.**
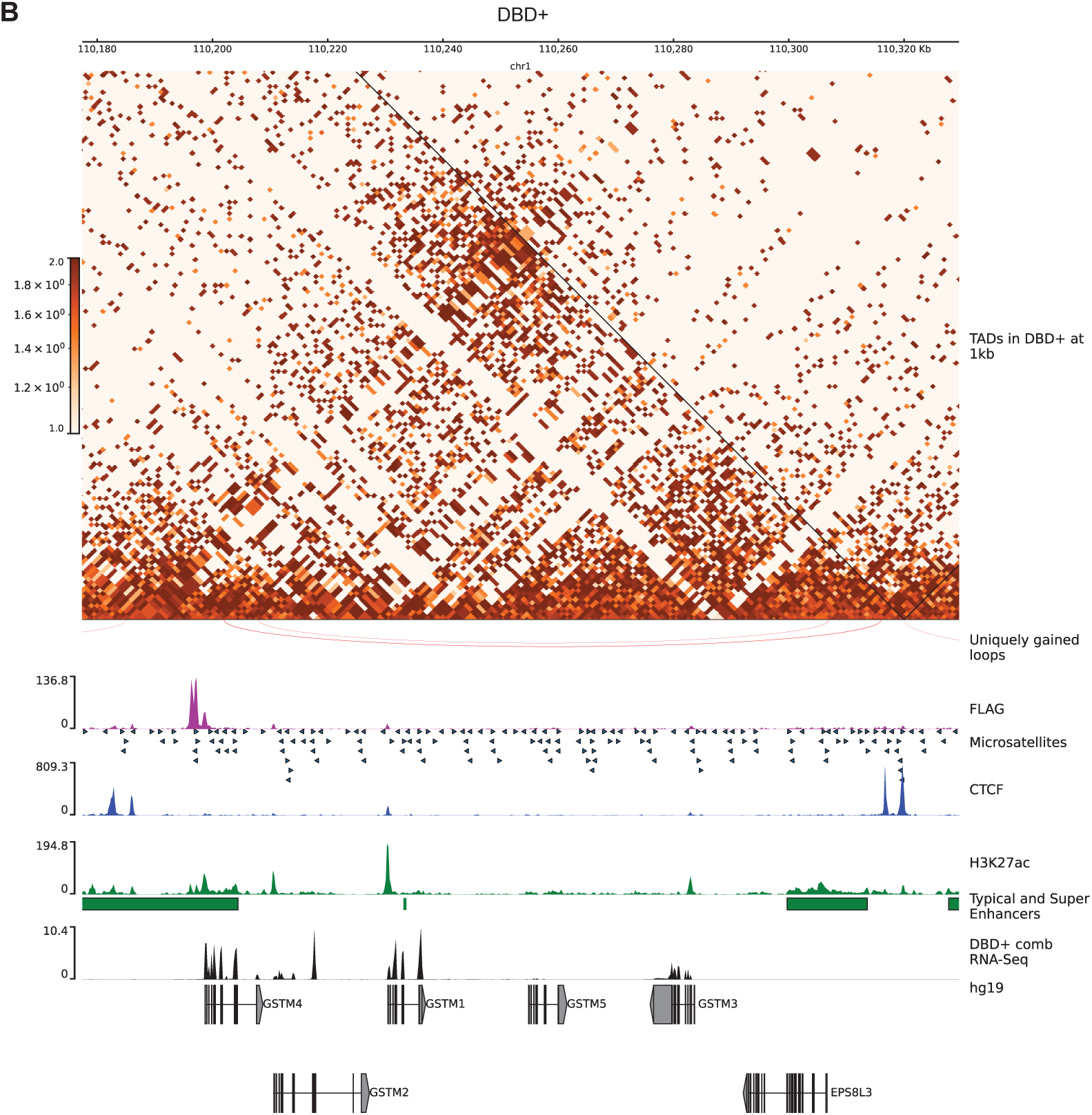
B. *GSTM4* hub chr 1 in A-673 DBD+ cells. TADs are depicted on 1kb matrices (DBD+/KD). Uniquely gained loops are shown as red inverted arcs. FLAG CUT&Tag bigwig tracks depicted in magenta. GGAA microsatellites in hg19. CTCF CUT&Tag track is in blue middle row. H3K27ac tracks are in green. Enhancers and super-enhancers are shown as green bars. Gene expression is in black tracks.

**Supplementary Figure 9.**
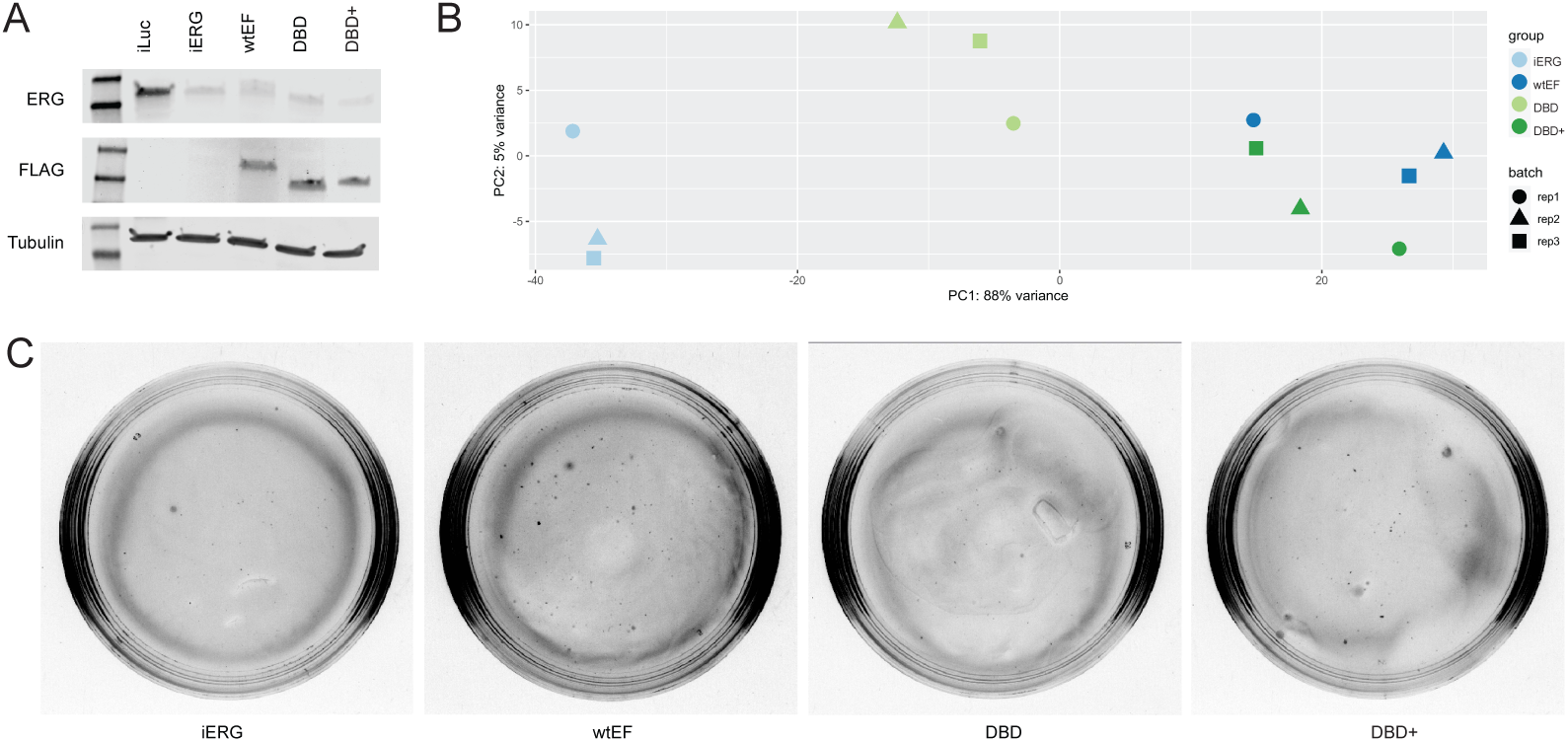
A. Knock-down of endogenous EWSR1::ERG detected with ERG ab and rescue of wtEF, DBD and DBD+ detected with FLAG ab. B. A PCA plot of RNA-seq experiments in TTC-466 cells. C. Representative images of soft agar colony plates.

**Supplementary Figure 10.**
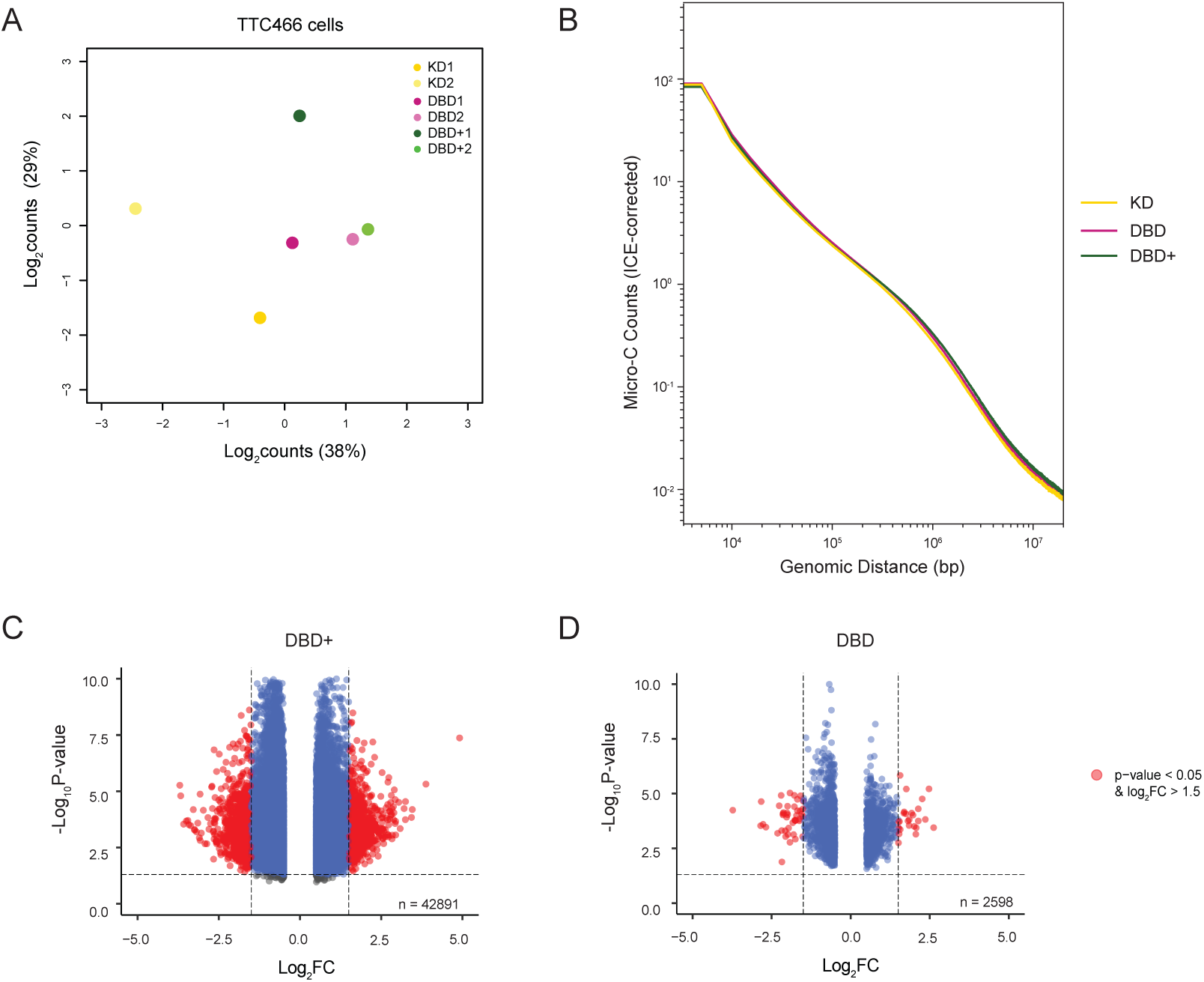
DBD-a4 helix of FLI1 domain is required to restructure chromatin in TTC-466 cell. A. Multidimensional scaling (MDS) plot of top 1000 interactions (500kb resolution) in each biological replicates. B. Genome-wide interaction frequency of combined replciates (ICE-corrected Micro-C counts) over genomic distance (bp) at 5kb resolution. C. Volcano plot showing differentially interacting regions (DIRs) detected at 500kb resolution for DBD+ replicates versus KD replicates. D. Volcano plot showing DIRs detected at same resolution for DBD replicates versus KD replicates.

**Supplementary Figure 11.**
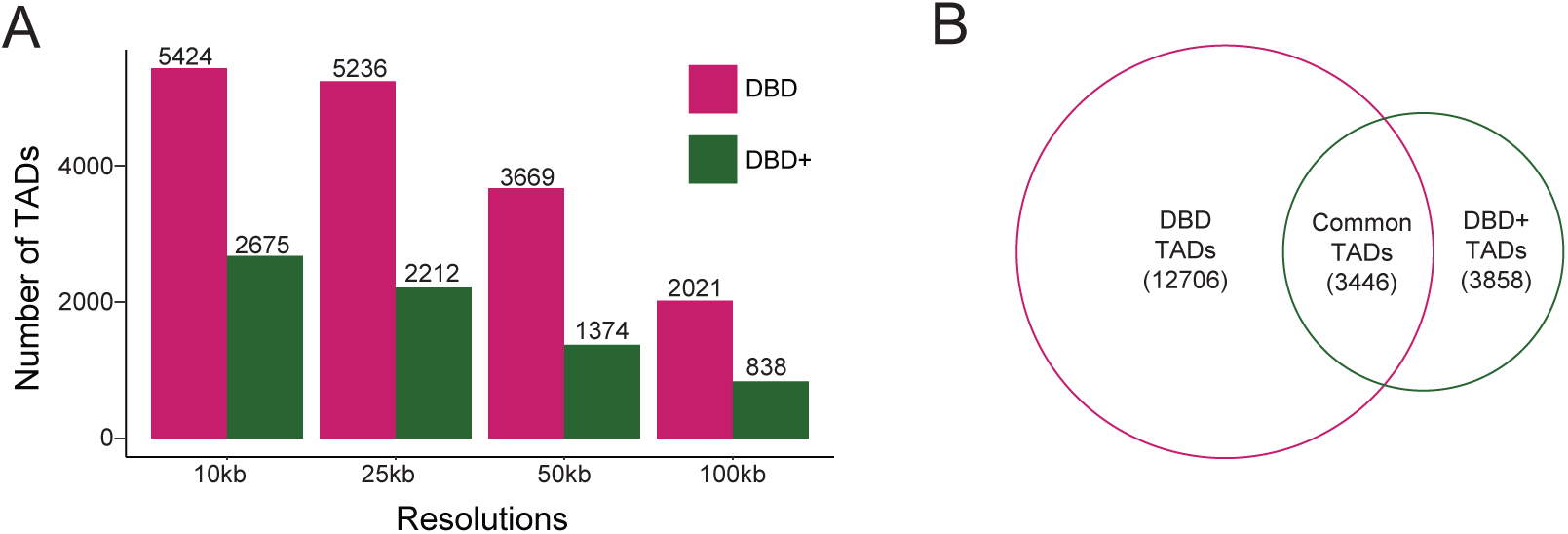
TAD analysis in TTC-466 cells. A. Number of TADs detected in DBD and DBD+ compared to KD at resolutions of 10kb, 25kb, 50kb and 100kb. B. Venn diagram of overlap between DBD and DBD+ TADs (compared to KD).

**Supplementary Figure 12.**
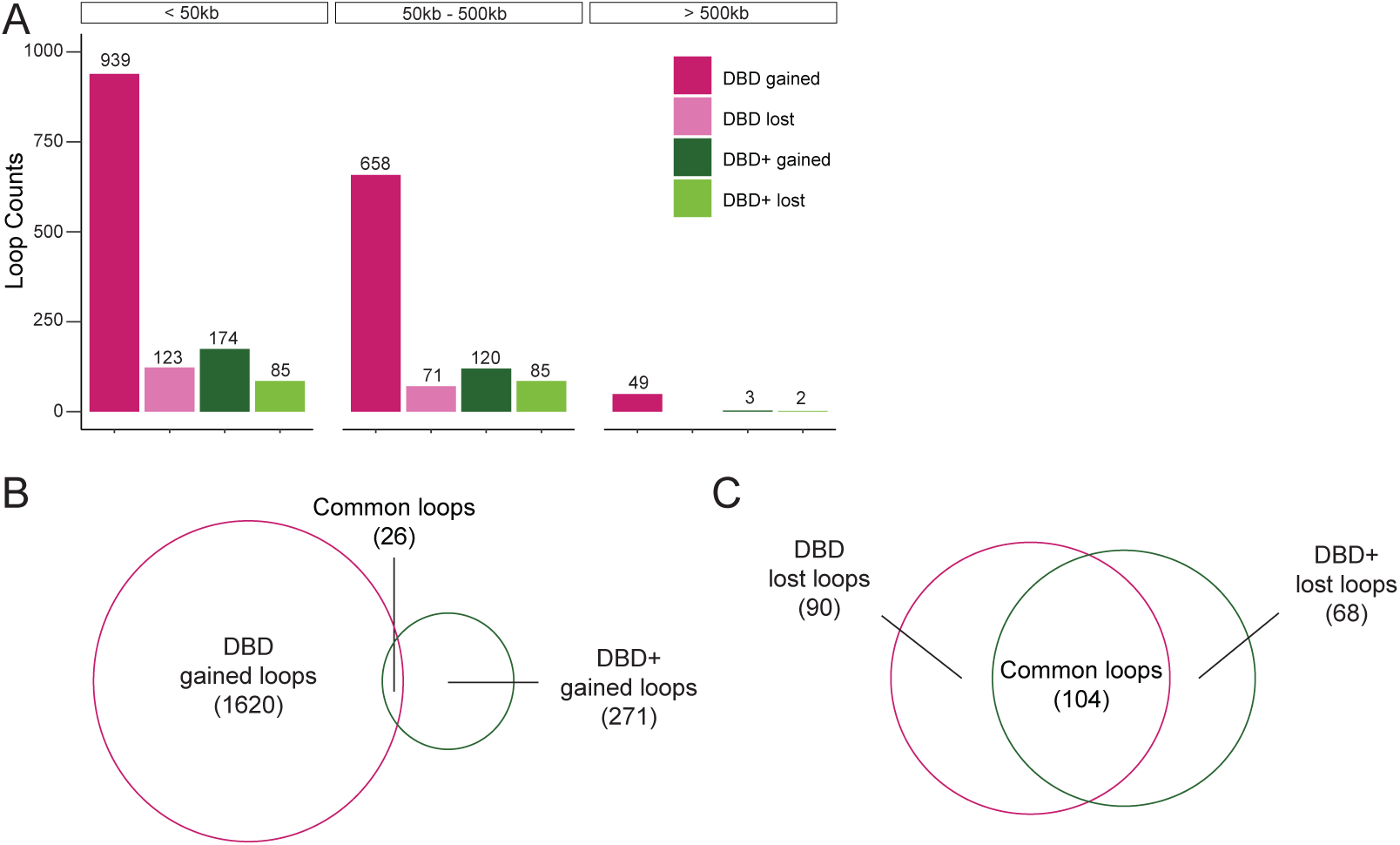
Loop analysis in TTC-466 cells. A. Number of loops detected in DBD and DBD+ compared to KD at resolutions of 1kb at short-range (<50kb), mid-range (50kb-500kb), and long-range (>500kb). B. Venn diagram of overlap between DBD and DBD+ uniquely gained loops (compared to KD). C. Venn diagram of overlap between DBD and DBD+ uniquely lost loops (compared to KD).

**Supplementary Figure 13.**
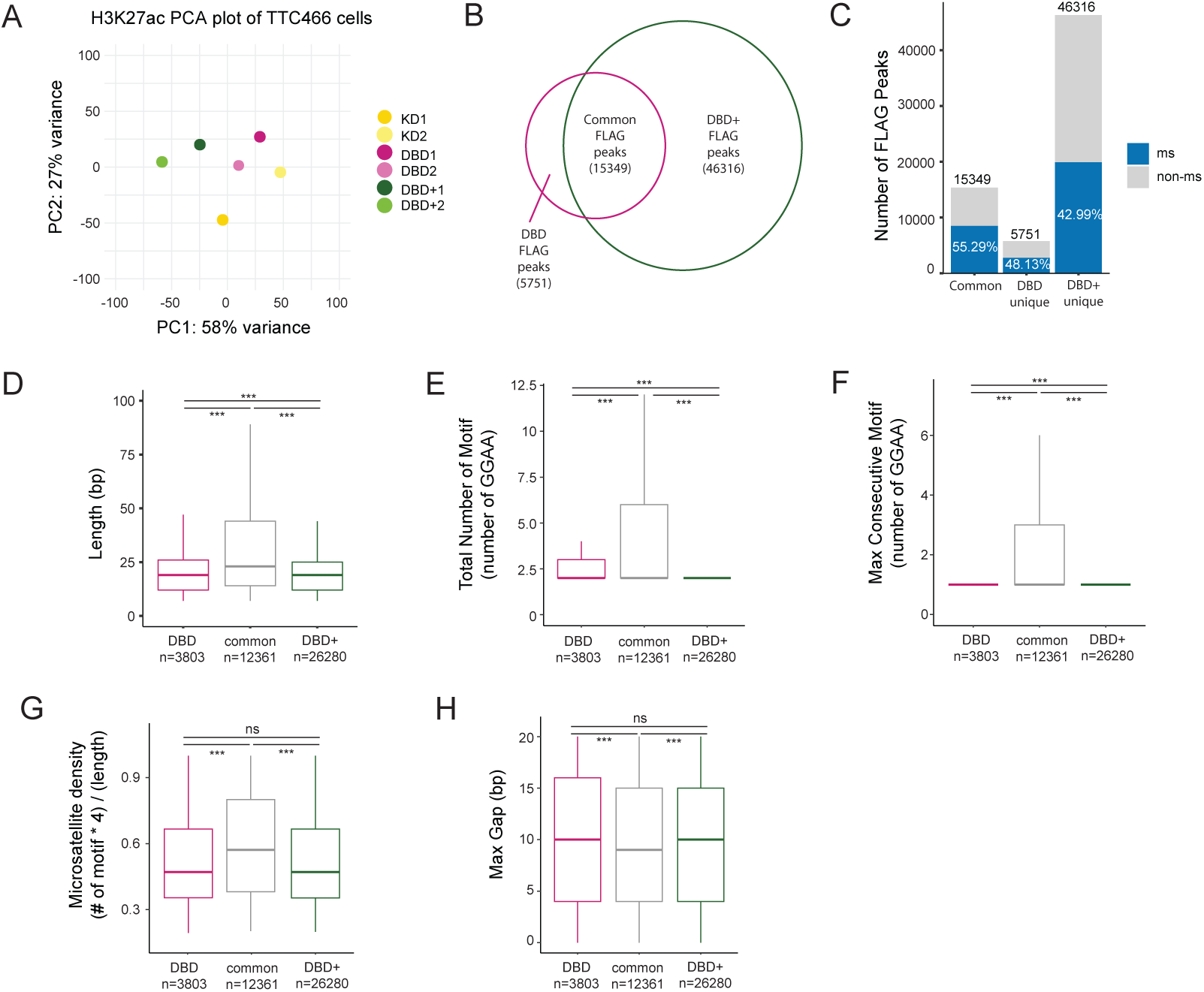
H3K27ac and FLAG CUT&Tag analysis in TTC-466 cells. A. PCA plot of H3K27ac peaks in biological replicates of KD, DBD and DBD+. B. Venn diagram of overlap between FLAG peaks (FDR < 0.05, FC > 8, counts > 80, IDR < 0.01) of DBD and DBD+ cells. C. Percentage of FLAG peaks bound at microsatellites in common, DBD unique and DBD+ unique peaks. D. Length (in bp) of GGAA microsatellites bound by DBD unique (mean=30.28), common in both (mean=37.79), and DBD+ unique (mean=22.74) FLAG peaks. E.Total number of GGAA motifs in microsatellites bound by DBD unique (mean=3.70), common in both (mean=6.05), and DBD+ unique (mean=3.00) FLAG peaks. F. Maximum consecutive number of GGAA motifs in microsatellites bound by DBD unique (mean=1.35), common in both (mean=3.44), and DBD+ unique (mean=1.57v) FLAG peaks G. Percent of GGAA motif in the microsatellites calculated as (# of motif x 4)/(length of microsatellites) bound by DBD unique (mean=0.53), common in both (mean=0.60), and DBD+ unique (mean=0.54) FLAG peaks. H. Maximum number of insertion (gaps in bp) in microsatellites bounds by DBD unique (mean=10.0), common (mean=9.36), and DBD+ unique (mean=9.77) FLAG peaks. * P value < 0.05, ** P value < 0.01, and *** P value < 0.001.

**Supplementary Figure 14.**
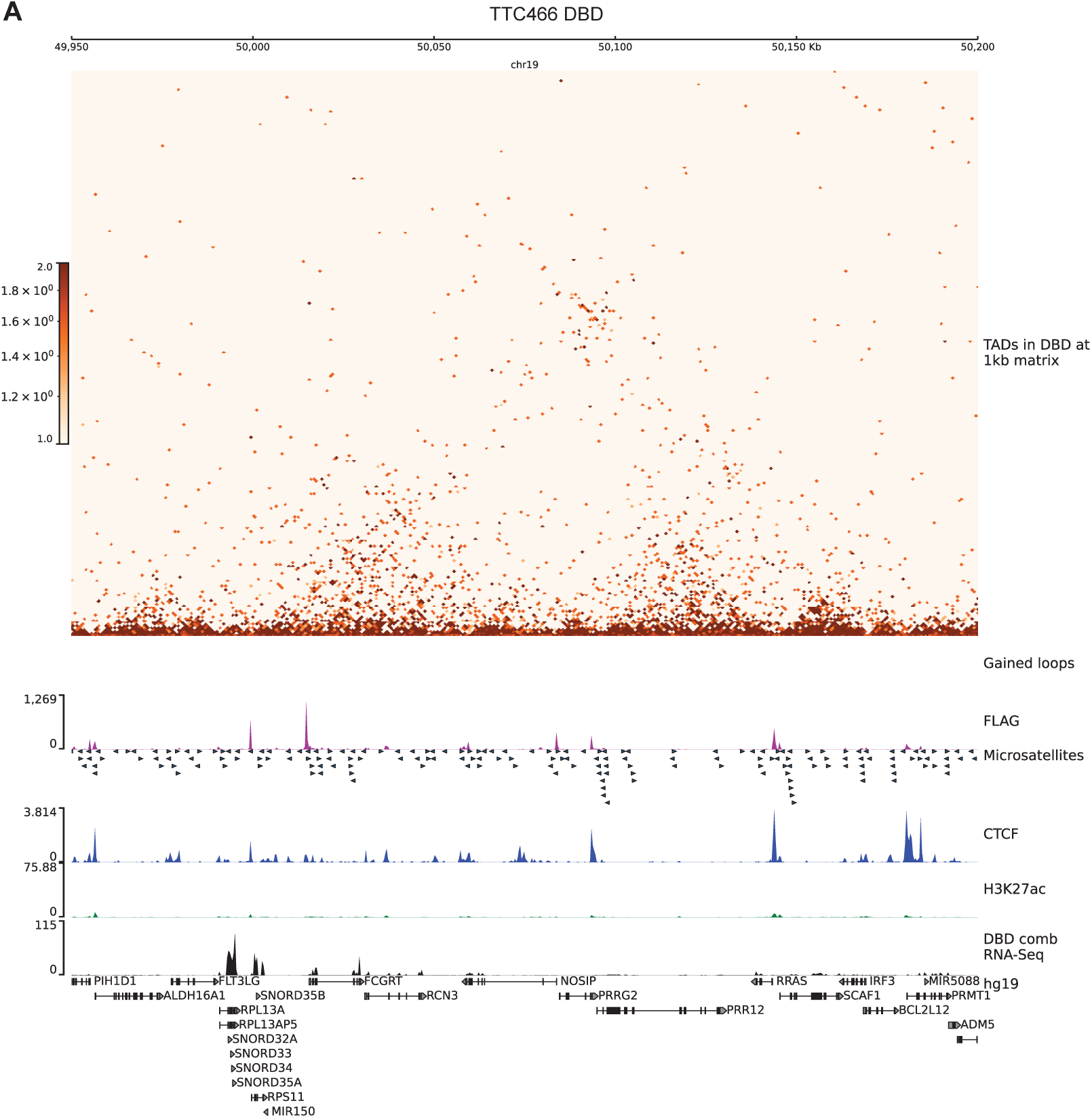
A. FCGRT hub in 250kb region on chr 19 in TTC-466 DBD cells. TADs are depicted on 1kb matrices (DBD/KD). Gained loops are shown as red inverted arcs. FLAG CUT&Tag bigwig tracks depicted in magenta. GGAA microsatellites in hg19. CTCF CUT&Tag track is in blue middle row. H3K27ac tracks are in green. Gene expression is in black tracks.

**Supplementary Figure 14.**
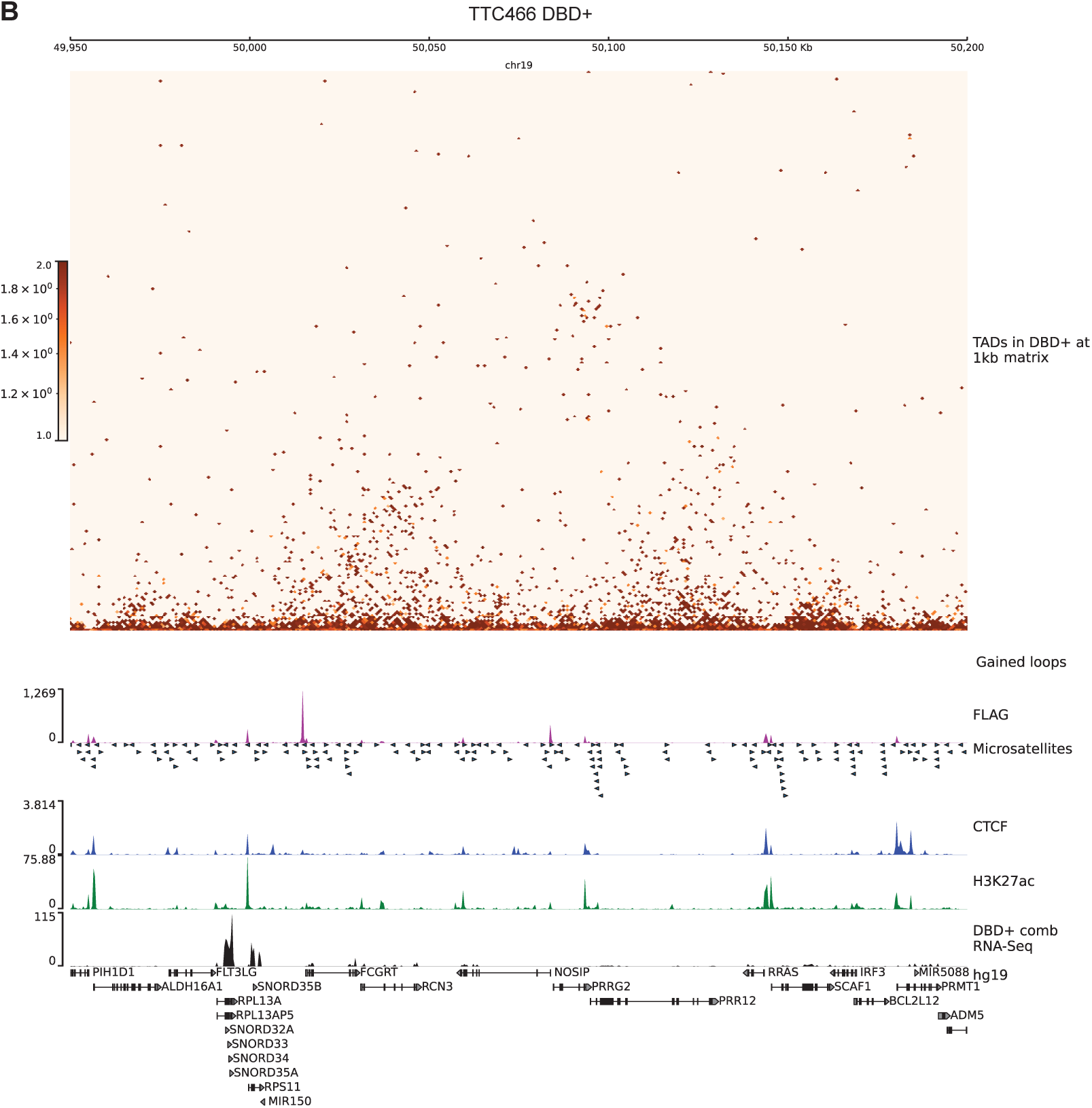
B. *FCGRT* hub in 250kb region on chr 19 in TTC-466 DBD+ cells. TADs are depicted on 1kb matrices (DBD+/KD). Gained loops are shown as red inverted arcs. FLAG CUT&Tag bigwig tracks depicted in magenta. GGAA microsatellites in hg19. CTCF CUT&Tag track is in blue middle row. H3K27ac tracks are in green. Gene expression is in black tracks.

**Supplementary Figure 15.**
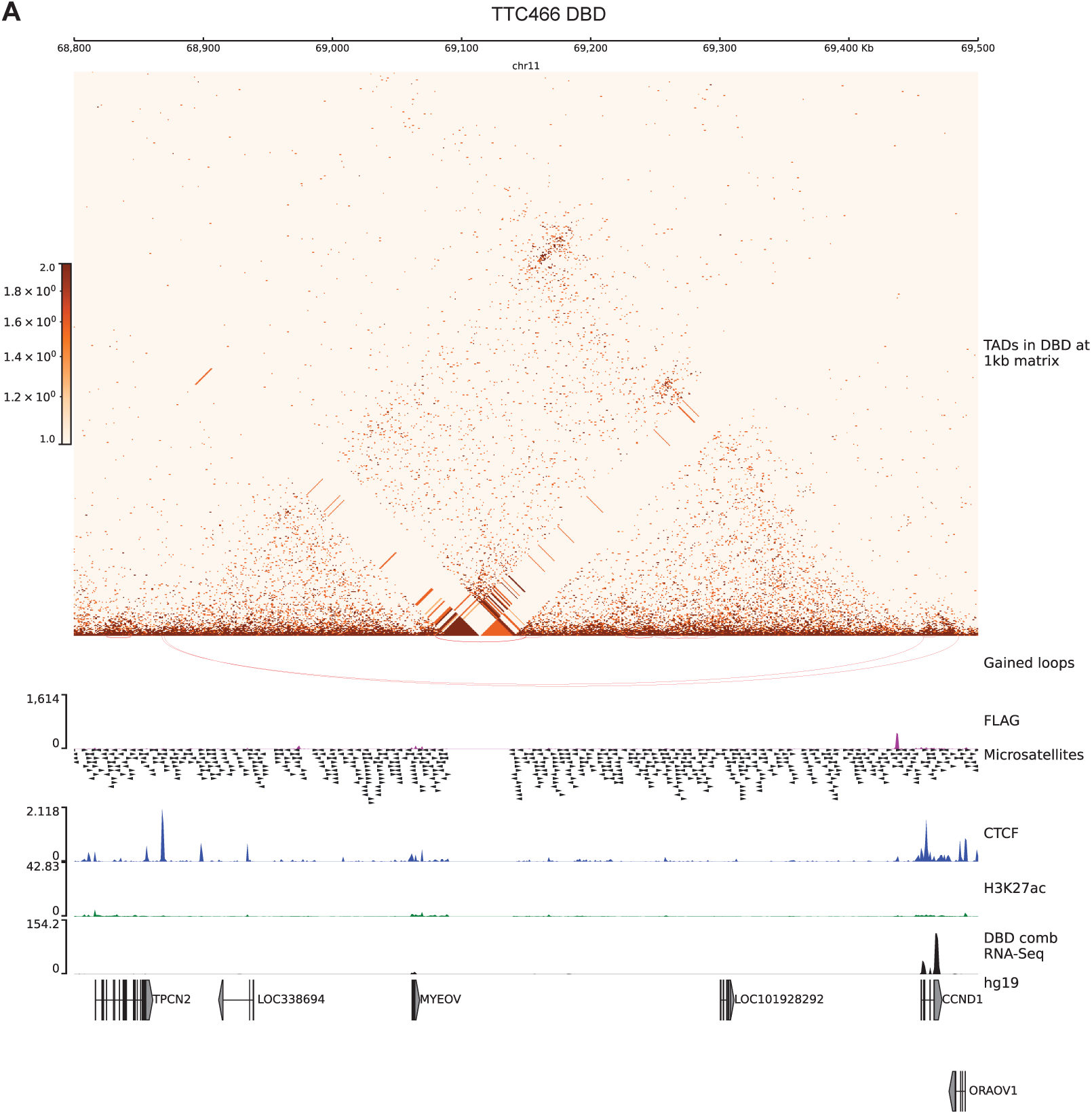
A. *CCND1* hub in 700kb region on chr 11 in TTC-466 DBD cells. TADs are depicted on 1kb matrices (DBD/KD). Gained loops are shown as red inverted arcs. FLAG CUT&Tag bigwig tracks depicted in magenta. GGAA microsatellites in hg19. CTCF CUT&Tag track is in blue middle row. H3K27ac tracks are in green. Gene expression is in black tracks.

**Supplementary Figure 15.**
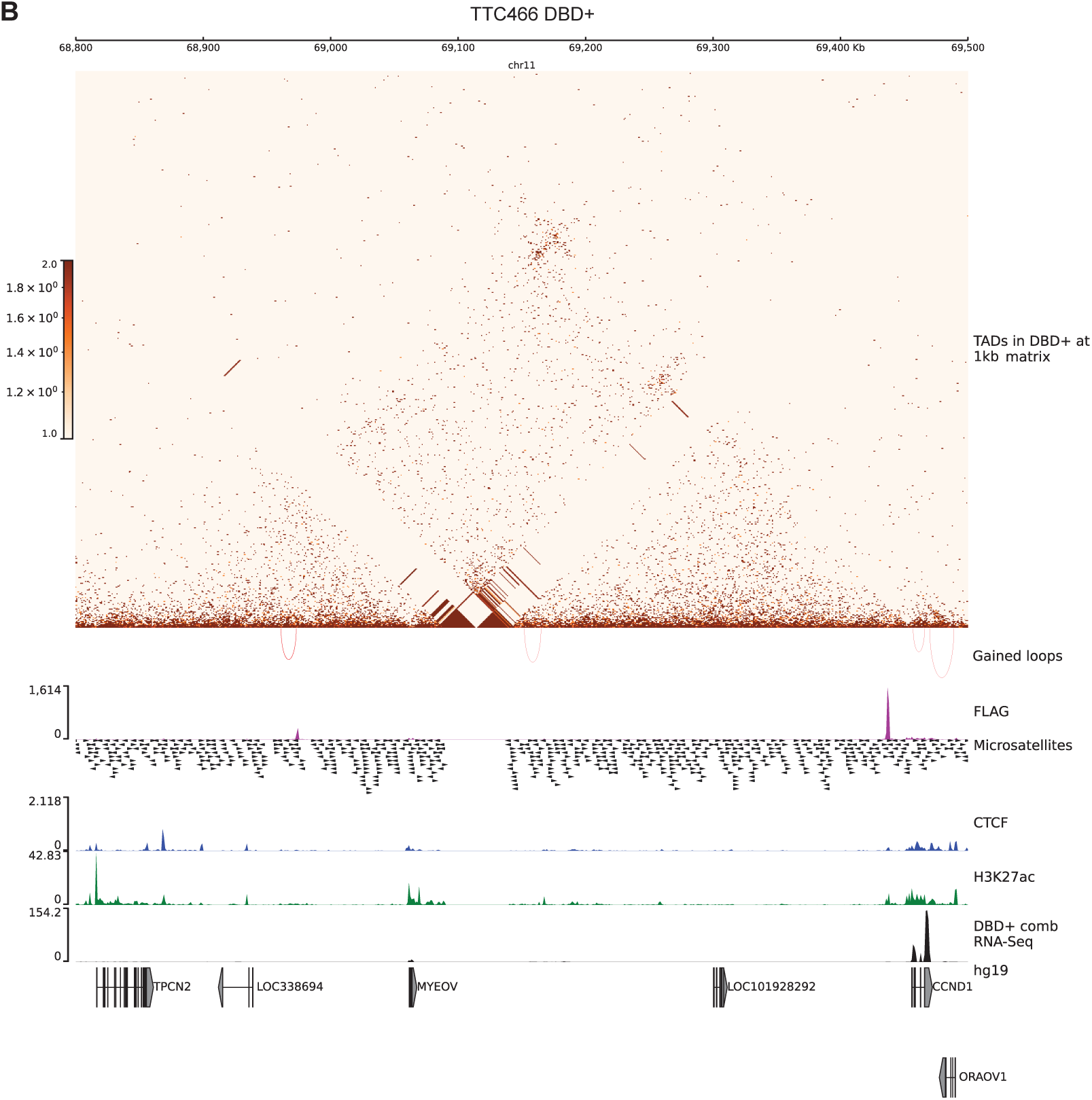
B. *CCND1* hub in 700kb region on chr 11 in TTC-466 DBD+ cells. TADs are depicted on 1kb matrices (DBD+/KD). Gained loops are shown as red inverted arcs. FLAG CUT&Tag bigwig tracks depicted in magenta. GGAA microsatellites in hg19. CTCF CUT&Tag track is in blue middle row. H3K27ac tracks are in green. Gene expression is in black tracks.

**Supplementary Figure 16.**
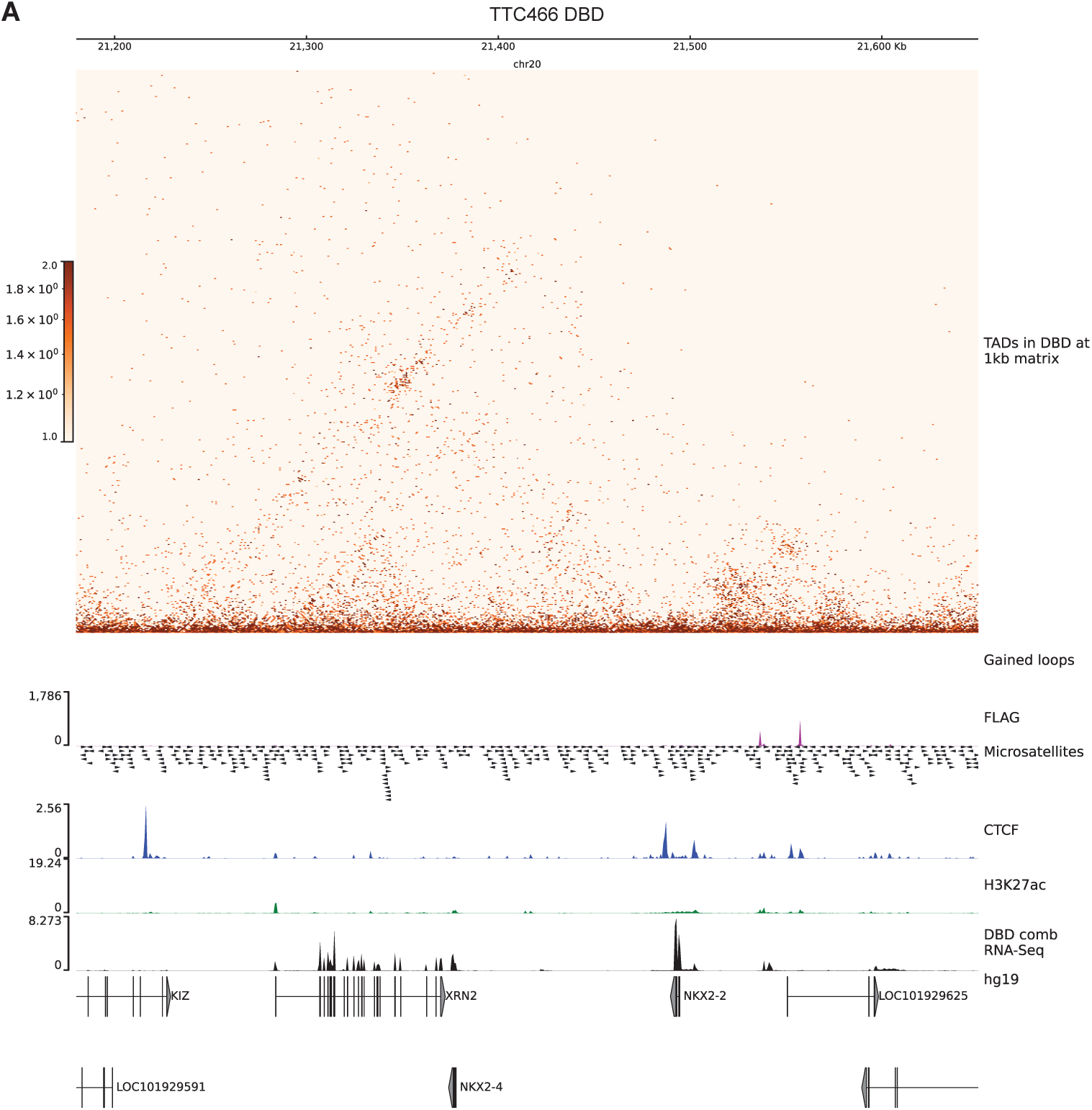
A. *NKX2-2* hub chr 20 in TTC-466 DBD cells. TADs are depicted on 1kb matrices (DBD/KD). Gained loops are shown as red inverted arcs. FLAG CUT&Tag bigwig tracks depicted in magenta. GGAA microsatellites in hg19. CTCF CUT&Tag track is in blue middle row. H3K27ac tracks are in green. Gene expression is in black tracks.

**Supplementary Figure 16.**
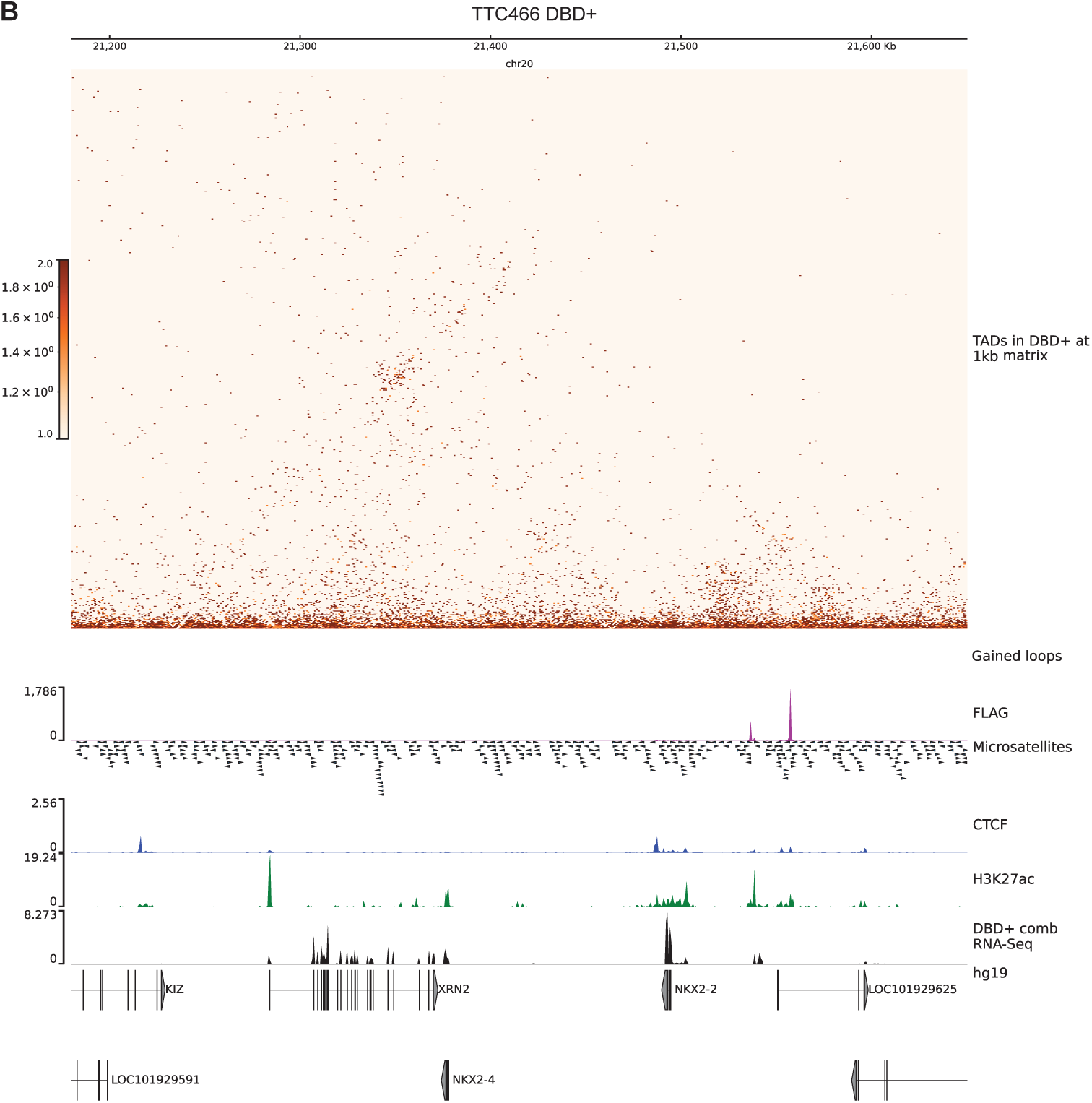
B. *NKX2-2* hub on chr 20 in TTC-466 DBD+ cells. TADs are depicted on 1kb matrices (DBD+/KD). Gained loops are shown as red inverted arcs. FLAG CUT&Tag bigwig tracks depicted in magenta. GGAA microsatellites in hg19. CTCF CUT&Tag track is in blue middle row. H3K27ac tracks are in green. Gene expression is in black tracks.

**Supplementary Figure 17.**
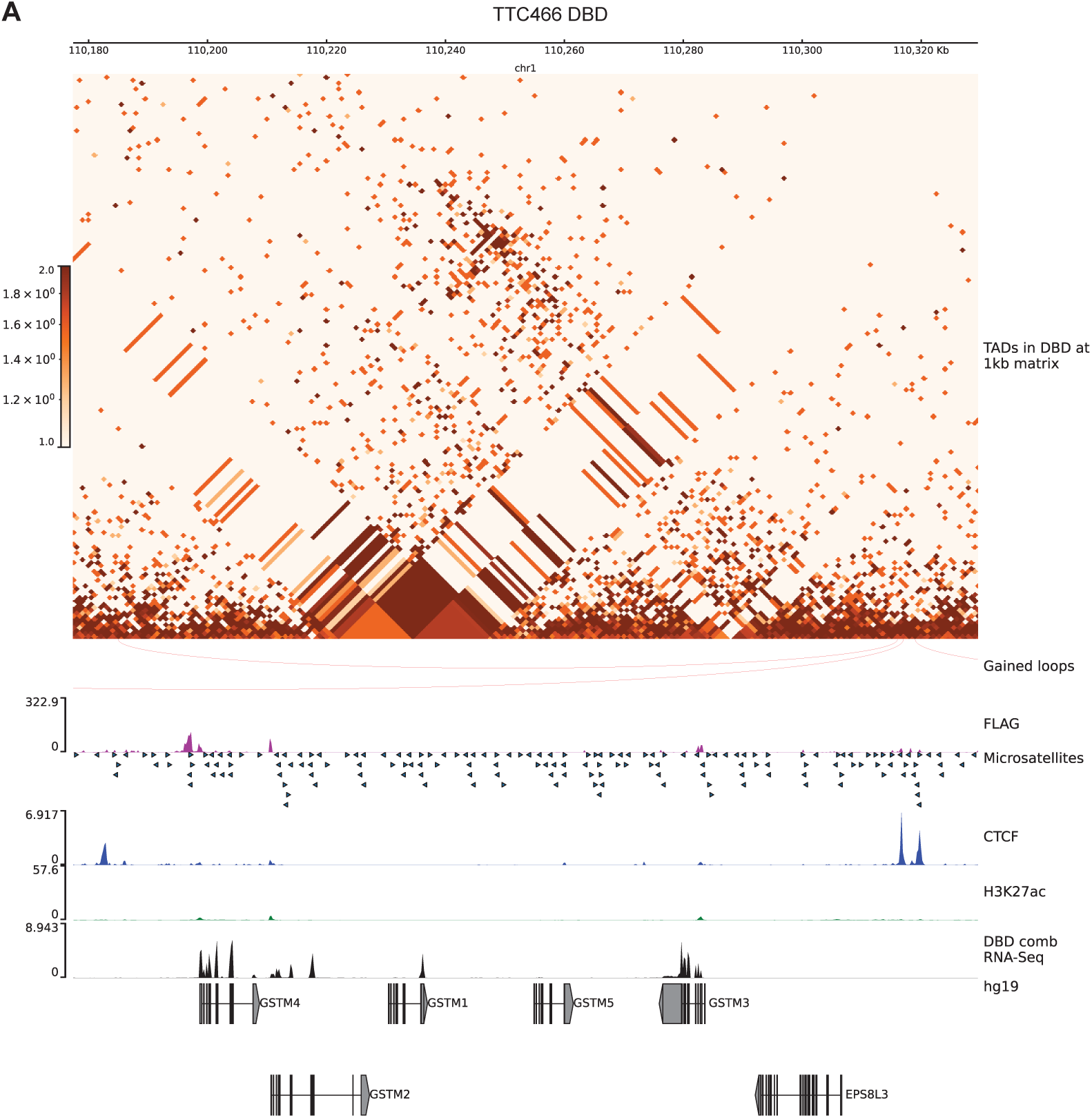
A. *GSTM4* hub chr 1in TTC-466 DBD cells. TADs are depicted on 1kb matrices (DBD/KD). Gained loops are shown as red inverted arcs. FLAG CUT&Tag bigwig tracks depicted in magenta. GGAA microsatellites in hg19. CTCF CUT&Tag track is in blue middle row. H3K27ac tracks are in green. Gene expression is in black tracks.

**Supplementary Figure 17.**
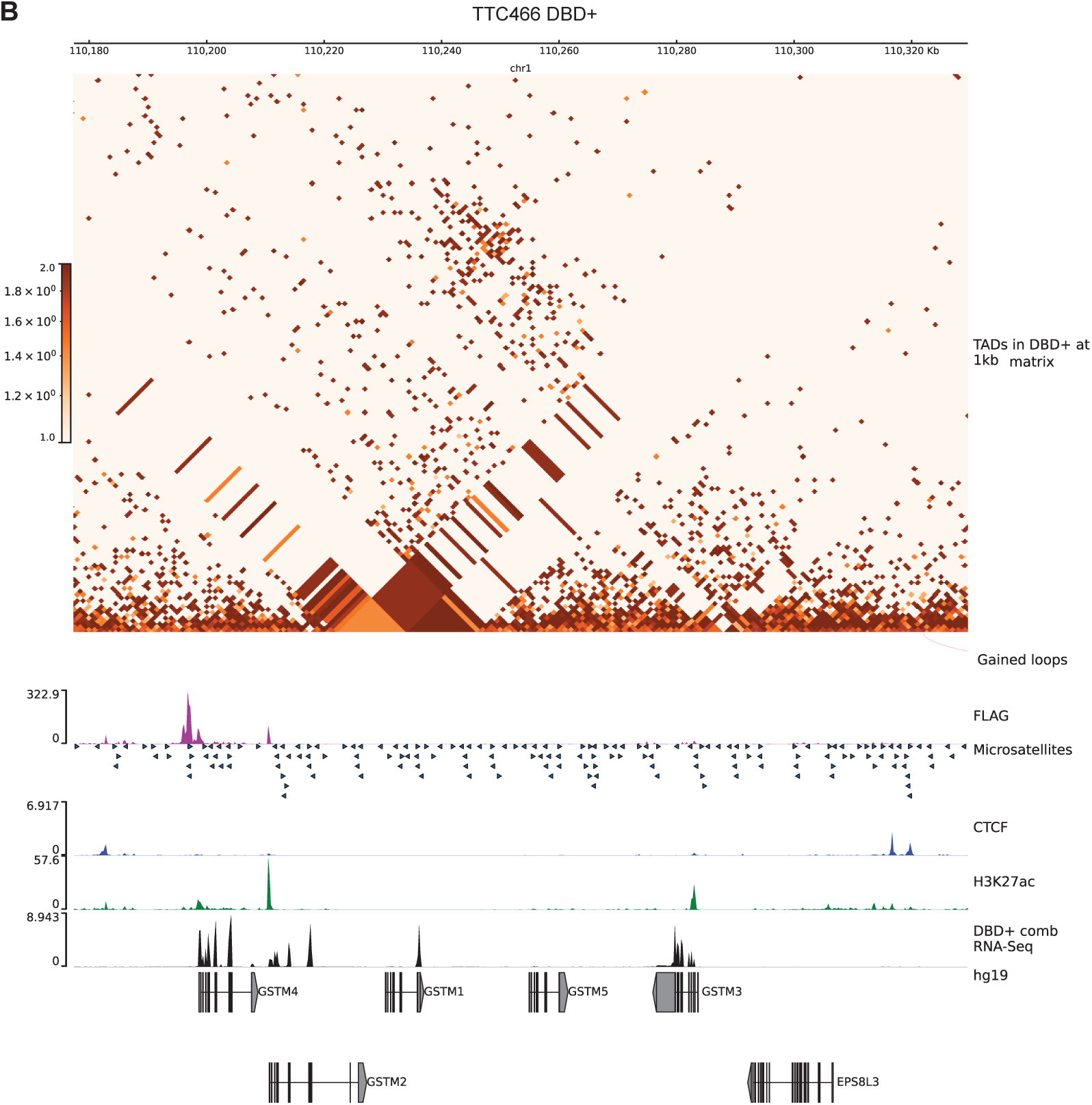
B. *GSTM4* hub on chr 1 in TTC-466 DBD+ cells. TADs are depicted on 1kb matrices (DBD+/KD). Gained loops are shown as red inverted arcs. FLAG CUT&Tag bigwig tracks depicted in magenta. GGAA microsatellites in hg19. CTCF CUT&Tag track is in blue middle row. H3K27ac tracks are in green. Gene expression is in black tracks.

**Supplementary Figure 18.**
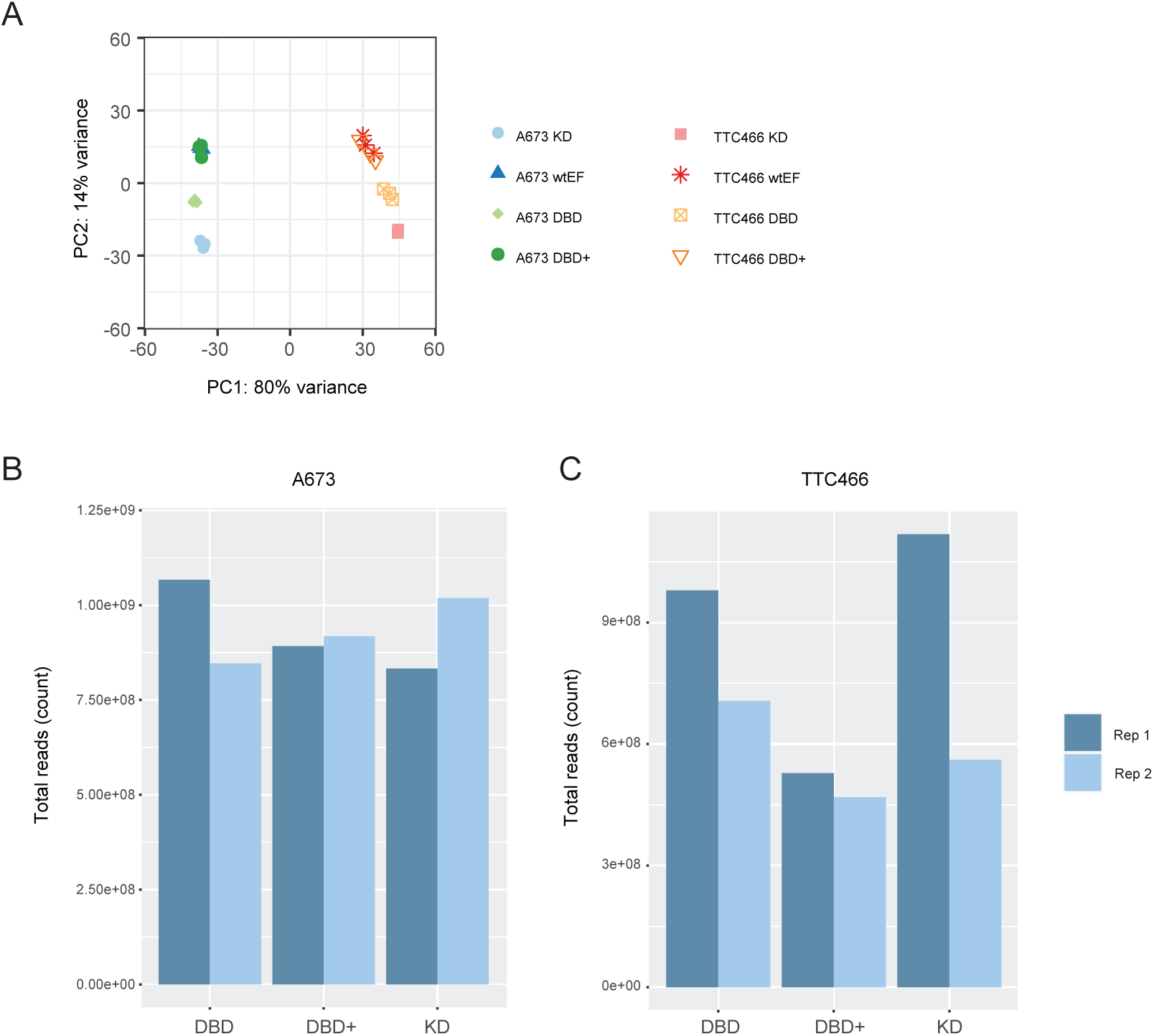
A-673 and TTC-466 cell line comparison. A. PCA plot of RNA-Seq replicates of A-673 and TTC-466 cells. B. Sequencing depth of Micro-C replicates of A-673 cells. C. Sequencing depth of Micro-C replicates of TTC-466 cells.

**Table 1.** Supplementary Table 1. Differential expression of *FCGRT* hub genes in DBD and DBD+ compared to KD.

| Gene Symbol | DBD FC | DBD padj | DBD+ FC | DBD+ padj |
| --- | --- | --- | --- | --- |
| *ALDH16A1* | 1.207 | 0.046 | 1.107 | 0.329 |
| *RPL13A* | -1.132 | 0.0214 | -1.209 | 7.46E-05 |
| *RPL13AP5* | -1.109 | 0.491 | -1.149 | 0.291 |
| *RPS11* | -1.068 | 0.337 | -1.177 | 0.002 |
| *FCGRT* | 1.639 | 2.14E-4 | 2.44 | 2.54E-13 |
| *RCN3* | -1.588 | 6.85E-06 | -1.406 | 9.58E-4 |
| *NOSIP* | 1.065 | 0.447 | -1.041 | 0.637 |
| *PRRG2* | 1.117 | 0.760 | -1.042 | 0.917 |
| *PRR12* | 1.177 | 0.007 | 1.088 | 0.204 |
| *RRAS* | -1.655 | 1.63E-17 | -1.383 | 5.10E-08 |
| *SCAF1* | 1.065 | 0.357 | -1.07 | 0.283 |
| *IRF3* | 1.081 | 0.402 | 1.196 | 0.0115 |
| *BCL2L12* | 1.255 | 0.007 | 1.359 | 6.60E-05 |
| *PRMT1* | 1.151 | 0.015 | 1.145 | 0.015 |
| *ADM5* | 1.122 | 0.740 | 1.064 | 0.864 |

## Data availability

The sequencing dataset generated and analyzed in this study are available in the Gene Expression Omnibus and accessible at GSE249578 for all A-673 data and GSE268935 (Micro-C), GSE268940 (FLAG), GSE268941 (H3K27ac), GSE268942 (CTCF), [GSE268944](https://www.ncbi.nlm.nih.gov/geo/query/acc.cgi?acc=GSE268942) (RNA-Seq) for TTC-466 cells. All other data not part of GEO submissions are available from the corresponding author on a reasonable request.

## Code availability

All codes used in analysis of the sequencing data are compiled in Supplementary files.

## Acknowledgements

We thank Dr. Timothy Triche at the Children’s Hospital of Los Angeles for generously providing us with TTC-466 cells. We also thank Ali Snedden, Yuan Zhang, and John Burian at High Performance Computing group at Nationwide Children’s Hospital for their support; and Institute for Genomic Medicine at Nationwide Children’s Hospital for sequencing support. We are grateful for the comments and discussion from members of Lessnick lab, Theisen lab and Bioinformatics group at Nationwide Children’s Hospital. We also thank Dr. Elaine R. Mardis for her thoughtful comments on this manuscript. This research was supported by NIH U54 CA231641 to S.L.L. and NIH T32 CA269052 to A.B.

